# Widespread gene-environment interactions shape the immune response to SARS-CoV-2 infection in hospitalized COVID-19 patients

**DOI:** 10.1101/2024.12.03.626676

**Authors:** Haley E Randolph, Raúl Aguirre-Gamboa, Elsa Brunet-Ratnasingham, Tomoko Nakanishi, Veronica Locher, Ellen Ketter, Cary Brandolino, Catherine Larochelle, Alexandre Prat, Nathalie Arbour, Anne Dumaine, Andrés Finzi, Madeleine Durand, J Brent Richards, Daniel E Kaufmann, Luis B Barreiro

## Abstract

Genome-wide association studies performed in patients with coronavirus disease 2019 (COVID-19) have uncovered various loci significantly associated with susceptibility to SARS-CoV-2 infection and COVID-19 disease severity. However, the underlying *cis*-regulatory genetic factors that contribute to heterogeneity in the response to SARS-CoV-2 infection and their impact on clinical phenotypes remain enigmatic. Here, we used single-cell RNA-sequencing to quantify genetic contributions to *cis*-regulatory variation in 361,119 peripheral blood mononuclear cells across 63 COVID-19 patients during acute infection, 39 samples collected in the convalescent phase, and 106 healthy controls. Expression quantitative trait loci (eQTL) mapping across cell types within each disease state group revealed thousands of *cis*-associated variants, of which hundreds were detected exclusively in immune cells derived from acute COVID-19 patients. Patient-specific genetic effects dissipated as infection resolved, suggesting that distinct gene regulatory networks are at play in the active infection state. Further, 17.2% of tested loci demonstrated significant cell state interactions with genotype, with pathways related to interferon responses and oxidative phosphorylation showing pronounced cell state-dependent variation, predominantly in CD14^+^ monocytes. Overall, we estimate that 25.6% of tested genes exhibit gene-environment interaction effects, highlighting the importance of environmental modifiers in the transcriptional regulation of the immune response to SARS-CoV-2. Our findings underscore the importance of expanding the study of regulatory variation to relevant cell types and disease contexts and argue for the existence of extensive gene-environment effects among patients responding to an infection.

## Main text

Susceptibility to viral infection varies widely among individuals, influenced by a combination of host genetics and environmental factors. However, the precise contribution of each to immune response variation and disease progression remains unclear. Recent advances have demonstrated the considerable role of host genetics in shaping human immune response variation through expression quantitative trait loci (eQTL) mapping, applied to various immune cell subsets both at baseline and after exposure to immune stimuli and live pathogens. These ‘immune response eQTL’ studies have identified numerous genetic variants that underlie differences in immune responses to infection, including both cell type-specific eQTL and eQTL induced only upon infection (i.e., response eQTL)^1–5^. However, a significant limitation of these studies is that immune response measurements were largely collected *in vitro*, raising questions about the role of gene-environment interactions during viral infection *in vivo*.

More recently, efforts have expanded to explore other forms of genetic interaction effects, facilitated by the availability of population-scale cohorts genotyped and characterized by single-cell RNA sequencing^6^. Continuous cell state-dependent eQTL—eQTL that interact with specific cellular contexts defined at single-cell resolution—have been shown to explain more variation in gene expression than conventional, non-interacting eQTL^7^. Notably, autoimmune risk variants were enriched in these state-dependent loci^7,8^, highlighting the critical importance of cellular context in understanding disease-relevant genetic variants.

The global COVID-19 pandemic highlighted the possible consequences of the spread of a novel virus in a naïve population. Particularly in the initial waves of the pandemic, substantial immune response variation and disease heterogeneity was observed among individuals infected with SARS-CoV-2, the virus that causes COVID-19. While a fraction of individuals succumbed to severe disease, some developed typical influenza-like symptoms, while others harbored asymptomatic SARS-CoV-2 infections^9^. Although much of this variation can be attributed to environmental and social determinants^10^, genetic factors also clearly play a role.

Genome-wide association studies (GWAS) conducted for SARS-CoV-2 susceptibility and COVID-19 severity phenotypes revealed a handful of genome-wide significant loci associated with these traits^11–13^, often in genes related to viral immunity, including *IFNAR2* and *OAS1*^13^. An eQTL mapping study performed in peripheral blood mononuclear cells (PBMCs) collected from healthy individuals exposed to SARS-CoV-2 *in vitro* also found that response eQTL were highly cell type-dependent, often specific to the SARS-CoV-2 infection condition in the myeloid compartment^5^. Despite these findings, few studies have examined how genome-wide *cis*-regulatory genetic variation influences immune response diversity directly in patients during active viral infection^14^.

In this study, we explore the nature of genetic interaction effects in the context of *bona fide* SARS-CoV-2 infection, using patient cells sampled prior to the rollout of COVID-19 vaccines and during longitudinal follow-up. We specifically investigate cell type-specific, disease state-specific, and cell state-dependent gene regulatory heterogeneity, providing new insights into how genetic variation shapes immune responses *in vivo*.

### Single-cell profiling reveals severity-dependent cellular restructuring in COVID-19 patients

In this study, we used single-cell RNA-sequencing to profile the transcriptomes of PBMCs collected from 106 healthy control donors, 63 hospitalized COVID-19 patients during the acute stage of infection (days after symptom onset [DSO] ≤ 20 days, mean DSO at time of sampling = 12.1 days), and 39 samples obtained from a subset of recovered COVID-19 patients resampled at various time points following their initial primary infection (“follow-ups”, DSO > 20 days, mean DSO = 128.8 days) (Fig. 1A, Fig. S1A, Table S1). Across individuals, we captured 361,119 high-quality single-cell transcriptomes (n = 163,639 cells from controls, n = 131,457 cells from acute patients, and n = 66,023 cells from follow-ups). Clustering followed by cell type label transfer annotation from a multimodal human PBMC reference dataset (Hao et al.^15^, detailed in Methods) revealed 30 distinct immune cell types at fine-scale resolution (Fig. 1B).

**Fig. 1.**
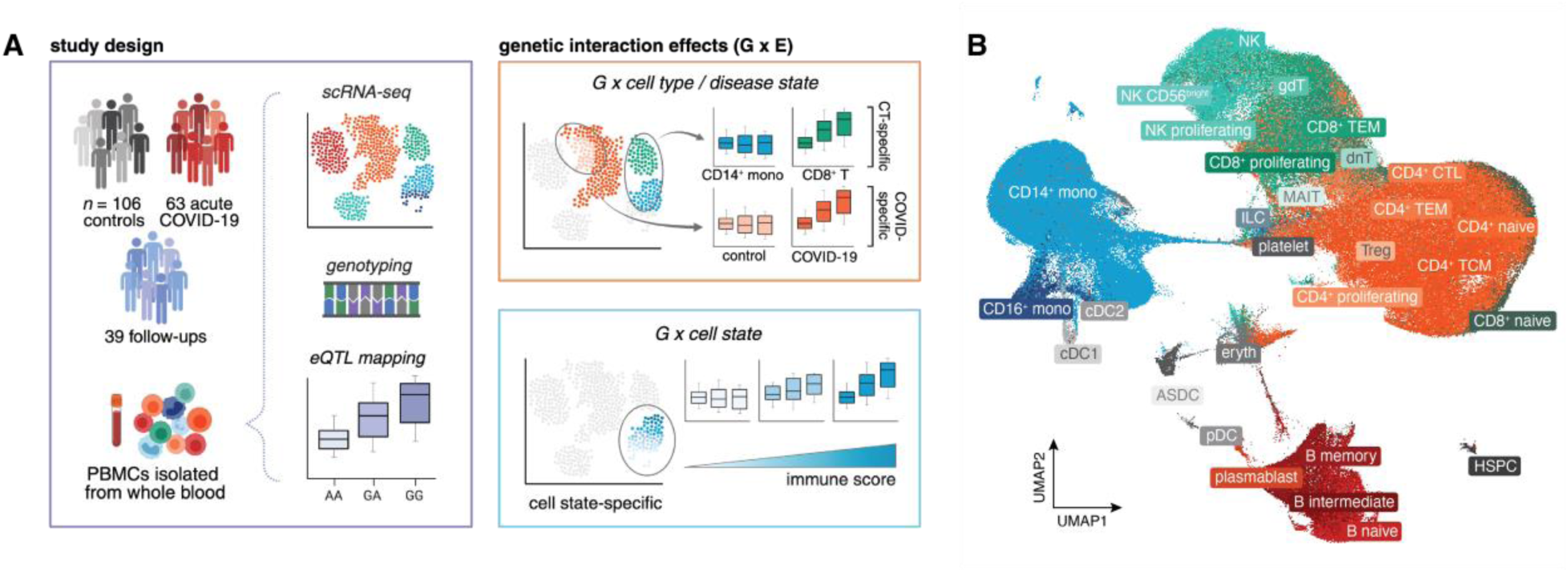
Summary of the study cohort and aims. **(A)** Study design (left) and examples of various gene-environment interactions, including cell type-, disease state-, and cell state-dependent effects, evaluated in this study (right). **(B)** UMAP visualization of all cells (n = 361,119) collected across healthy control, acute COVID-19 patient, and follow-up samples (n = 208 samples). ASDC: AXL^+^SIGLEC6^+^ dendritic cells, CD4^+^ CTL: cytotoxic CD4^+^ T cells, cDC: conventional dendritic cells, dnT: double-negative T cells, Eryth: erythrocytes, gdT: gamma delta T cells, HSPC: hematopoietic stem and progenitor cells, ILC: innate lymphoid cells, MAIT: mucosal associated invariant T cells, mono: monocytes, NK: natural killer, pDC: plasmacytoid dendritic cells, TEM: T effector memory, TCM: T central memory.

We next sought to dissect the extent to which SARS-CoV-2 infection induces shifts in underlying cell type composition across acutely-infected individuals compared to non-infected healthy controls and recovered donors. Although all COVID-19 patients included in this study were hospitalized at the time of sample collection, these patients spanned a range of clinical disease severity, allowing us to evaluate the effect of severity on various molecular phenotypes. Disease severity was assessed using a five-point scale of respiratory support needed at the time of acute patient sampling, encompassing the following categories: Moderate (MOD, n = 16), Severe (SEV, n = 17), 2-Critical (CRIT2, n = 9), 3-Critical (CRIT3, n = 20), and 4-Critical (CRIT4, n = 1). A summary of basic demographic information stratified by disease severity can be found in Table S1. Non-critical patients were defined as those requiring no oxygen supplementation (moderate disease) or oxygen supplementation through a nasal cannula (severe disease), whereas critical patients required mechanical ventilation, ranging from non-invasive ventilation (CRIT2) and intubation (CRIT3) to extracorporeal membrane oxygenation (CRIT4).

We found that SARS-CoV-2 infection remodels the baseline cell type composition of PBMCs observed in healthy individuals, with the magnitude of disease severity further modifying this effect. The myeloid compartment displayed the most obvious infection- and severity-dependent changes: classical CD14^+^ monocytes were markedly expanded in all patient groups compared to healthy donors (p < 1 x 10^-10^ for all comparisons against controls; here, all critical patients [CRIT2 – 4] were considered as a single group), with the greatest expansion seen in severe and critical cases (Fig. 2A). In the follow-up samples, CD14^+^ monocyte proportions reverted back to frequencies similar to those seen in baseline healthy control donors (Fig. 2A), suggesting that this monocytic expansion is indeed infection-induced. Further, we observed that the frequency of CD14^+^ monocytes was strongly associated with disease severity, with more severe cases consistently displaying a greater proportion of classical monocytes (Pearson’s r = 0.60, p = 6.8 x 10^-6^) (Fig. 2B).

**Fig. 2.**
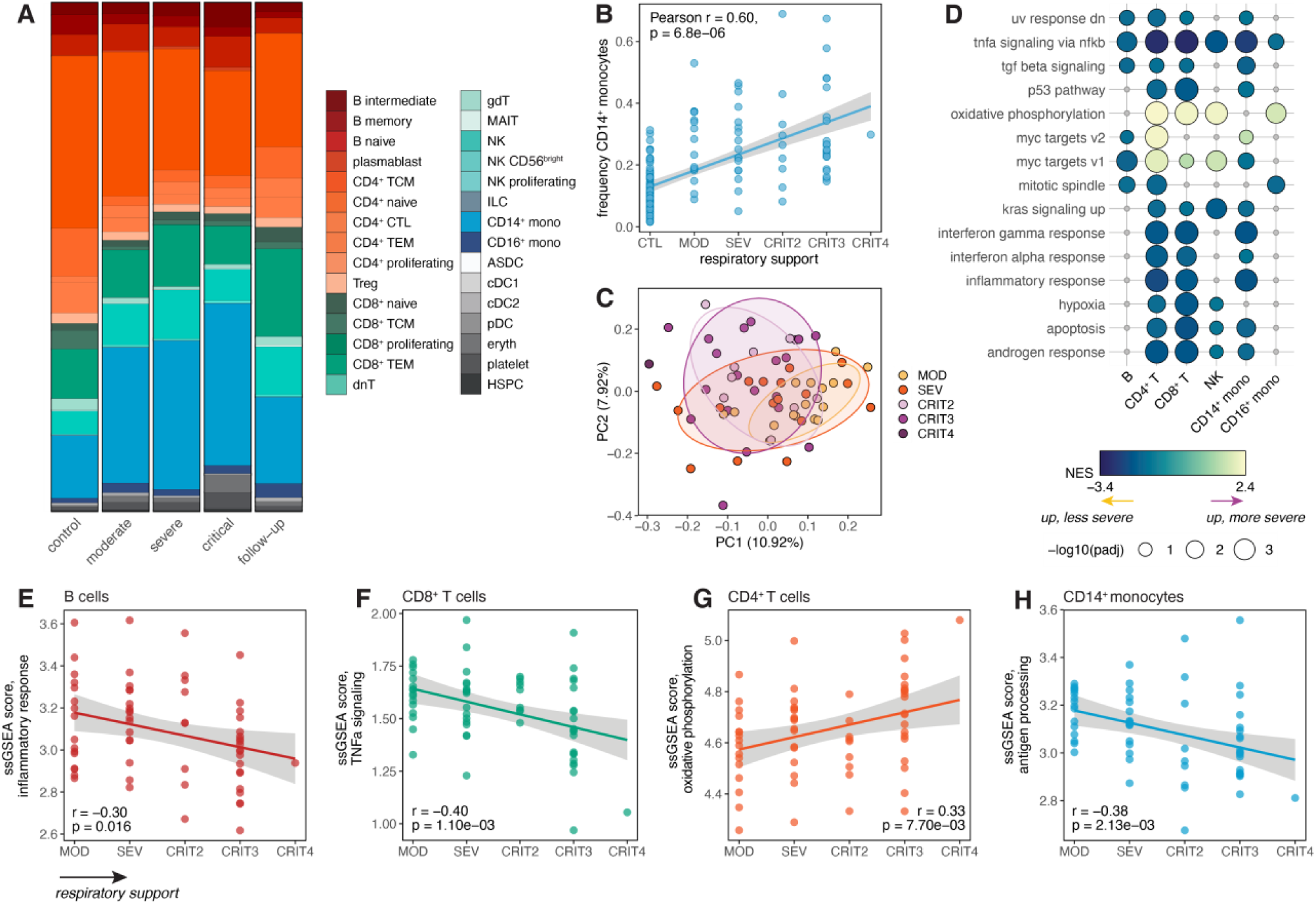
Effects of COVID-19 disease severity on underlying cell type composition and transcriptional signatures in hospitalized patients. **(A)** Cell type proportions stratified by disease severity at the time of sample collection. **(B)** Correlation between respiratory support score at the time of patient sampling and frequency of CD14^+^ monocytes. **(C)** PCA decomposition of the CD14^+^ monocyte expression data in COVID-19 patients colored by respiratory support score. **(D)** Hallmark enrichments for severity effects in COVID-19 patients across cell types. Colored circles represent pathways with FDR < 0.10; gray circles represent non-significant pathways. Only pathways significant in three or more cell types are shown. **(E-H)** Correlation between respiratory support score and ssGSEA scores in various cell types for **(E)** inflammatory response, **(F)** TNF-α signaling, **(G)** oxidative phosphorylation, and **(H)** antigen processing. In **(B)** and **(E-H)**, p-values and best-fit lines were obtained from linear regression models.

We also detected reductions of CD56^bright^ natural killer (NK) cells (p < 2 x 10^-4^) and plasmacytoid dendritic cells (pDC) (p < 4 x 10^-3^) in all severity groups compared to non-infected individuals (Fig. 2A). pDCs are known for their ability to secrete large quantities of type I interferon (IFN) following viral infection^16^, and NK cells are key facilitators of antiviral immunity, with CD56^bright^ NK cells being efficient producers of IFN-γ, TNF-α, and GM-CSF^17^. Our observations are in line with previous studies showing reductions in frequencies of both NK cells and pDCs in critical patients compared to healthy controls^18–21^. Together, this suggests that SARS-CoV-2 infection induces atypical cell type composition that largely resolves after the infection clears, particularly in cell populations known to be important in cytokine production and antiviral immune responses.

### Disease severity underlies variation in the transcriptional response to SARS-CoV-2 in hospitalized COVID-19 patients

To tease apart how variation in disease severity influences the transcriptional immune response to SARS-CoV-2 across cell types, we formally modeled the effect of severity on global gene expression estimates among COVID-19 patients sampled during the acute phase of disease (n = 63) within each cell type independently. In these analyses, we defined a set of top-level cell type populations by combining our fine-scale clusters into major groups corresponding to the six main cell types that comprise PBMCs, including CD4^+^ T cells, CD8^+^ T cells, B cells, NK cells, CD14^+^ monocytes, and CD16^+^ monocytes. Within this broader set of cell populations, we collapsed our single-cell gene expression estimates into pseudobulk estimates per sample, generating six bulk-like gene expression matrices that were used for subsequent modeling. We considered respiratory support score (described above) as a proxy for overall disease severity, and modeled severity score as a numeric variable, which allowed us to capture genes with expression levels linearly correlated with severity.

By far, CD14^+^ monocytes showed the largest number of genes associated with severity (n = 1,613, 14.8% of the transcriptome; FDR < 0.05), while other cell types had much less prominent effects (< 1.0% severity-associated genes) (Table S2). As expected, severity-associated genes largely overlapped those distinguishing COVID-19 patients from healthy controls (i.e., infection-associated genes, |log_2_FC| > 0.5, FDR < 0.05) across cell types (gene set overlap: 2.1-fold, p < 1 x 10^-10^) (Fig. S1B, S1C, Table S3). Principal component analysis (PCA) on the CD14^+^ monocyte pseudobulk expression data revealed that variation in disease severity had a noticeable impact on the transcriptional response of these cells, reflected in principal component (PC) 1 (10.9% percent variance explained [PVE]) and PC2 (7.9% PVE), which both separated non-critical patients (moderate/severe) from critical patients (Fig. 2C).

We then performed gene set enrichment analysis for the MSigDB Hallmark pathways^22^ to define the functional pathways differentiating the transcriptional signatures of COVID-19 patients along the spectrum of disease severity in our cohort (Fig. 2D, Table S4). We identified various immune response pathways significantly associated with severity, including TNF-α signaling via NF-κB in all cell types tested (FDR = 0.03 in CD16^+^ monocytes and FDR < 2 x 10^-3^ in other cell types), and IFN-γ response (FDR < 4 x 10^-4^), IFN-α response (FDR < 0.08), and inflammatory response (FDR < 4 x 10^-4^) in CD14^+^ monocytes, CD4^+^ T cells, and CD8^+^ T cells (Fig. 2D). All of these enrichments were detected among genes more highly expressed in less severe cases, suggesting that such patients engage stronger proinflammatory and antiviral immune responses compared to those with more severe disease presentations. Importantly, these findings are unlikely to be confounded by potential sampling biases, as sampling time point (i.e., DSO) showed no significant association with respiratory support score (Pearson r = 0.11, p = 0.37) (Fig. S1D). With the exception of TNF-α signaling, these pathway enrichments were cell type-specific, implicating classical monocytes, helper T cells, and cytotoxic T cells as the subsets most influenced by variation in disease severity and morbidity. Only the oxidative phosphorylation pathway was consistently elevated in more severe cases (FDR < 1.5 x 10^-3^ in CD4^+^ T cells, CD8^+^ T cells, NK cells, and CD16^+^ monocytes), suggesting a rewiring of metabolism in patients who poorly respond to SARS-CoV-2 (Fig. 2D).

To better characterize severity-associated heterogeneity in the transcriptional immune response, we computed single-sample gene set enrichment analysis (ssGSEA) scores capturing the activity of various functional pathways within each sample across cell types (detailed in Methods). Consistent with our enrichment analyses, the level of respiratory support was negatively correlated with ssGSEA inflammatory response scores (Pearson r = −0.30, p = 0.016) (Fig. 2E) and TNF-α signaling scores (Pearson r = −0.40, p = 1.1 x 10^-3^) (Fig. 2F) in B cells and CD8^+^ T cells, respectively. Similarly, respiratory support score was also positively associated with oxidative phosphorylation scores in CD4^+^ T cells (Pearson r = 0.33, p = 7.7 x 10^-3^) (Fig. 2G). Moreover, we created an antigen processing and presentation score based on the corresponding Biological Process gene set^23^, given the previously reported finding that SARS-CoV-2 inhibits the major histocompatibility complex (MHC) class I pathway, a pathway that plays a crucial role in antiviral immunity in lung epithelial cells^24^. Antigen processing scores were negatively correlated with severity in CD14^+^ monocytes (Pearson r = −0.38, p = 2.1 x 10^-3^) (Fig. 2H), while no significant association was found in any other cell type that we tested (p > 0.20), indicating that antigen presentation-associated functions are shut down in circulating classical monocytes in severe patients.

### Genetic interaction effects shape transcriptional response variation during acute SARS-CoV-2 infection

All individuals were genotyped for 4.19 million single nucleotide polymorphisms (SNPs), allowing us to delineate the role of *cis*-regulatory genetic and gene-environment interaction effects in the context of SARS-CoV-2 infection in patient-derived cells. To directly measure the contribution of cell type-specific and disease state-specific genetic variation during the course of a viral infection, we mapped *cis*-eQTL, defined as SNPs located either within or flanking (±100 kilobases, kb) each gene of interest, using the pseudobulk expression estimates for all six major cell types independently in i) healthy controls and ii) COVID-19 patients sampled during acute infection. To increase our power to detect shared and cell type- or disease state-specific effects, we utilized a multivariate adaptive shrinkage framework (mash)^25^ to leverage information about the underlying correlation structure within our dataset.

Across cell types and infection conditions, we identified 2,725 genes with at least one significant *cis*-eQTL [local false sign rate (lfsr) < 0.10 in at least one cell type-condition pair, 35.6% of genes tested; referred to as eGenes] (Fig. 3A, Table S5). B cells (n eGenes = 1,481) and CD16^+^ monocytes (n eGenes = 1,438) exhibited the fewest genetic effects, while CD14^+^ monocytes displayed the greatest number (n eGenes = 2,127) (Fig. 3A). Most genetic effects were shared between healthy individuals and COVID-19 patients within a given cell type—84.8% on average, referred to as ‘shared’ eGenes (lfsr_CTL_ < 0.1 and lfsr_COVID_ < 0.3 or vice versa)—and many of these shared eGenes were also common across cell types, with 59.0% shared across four or more cell types (Fig. S2).

**Fig. 3.**
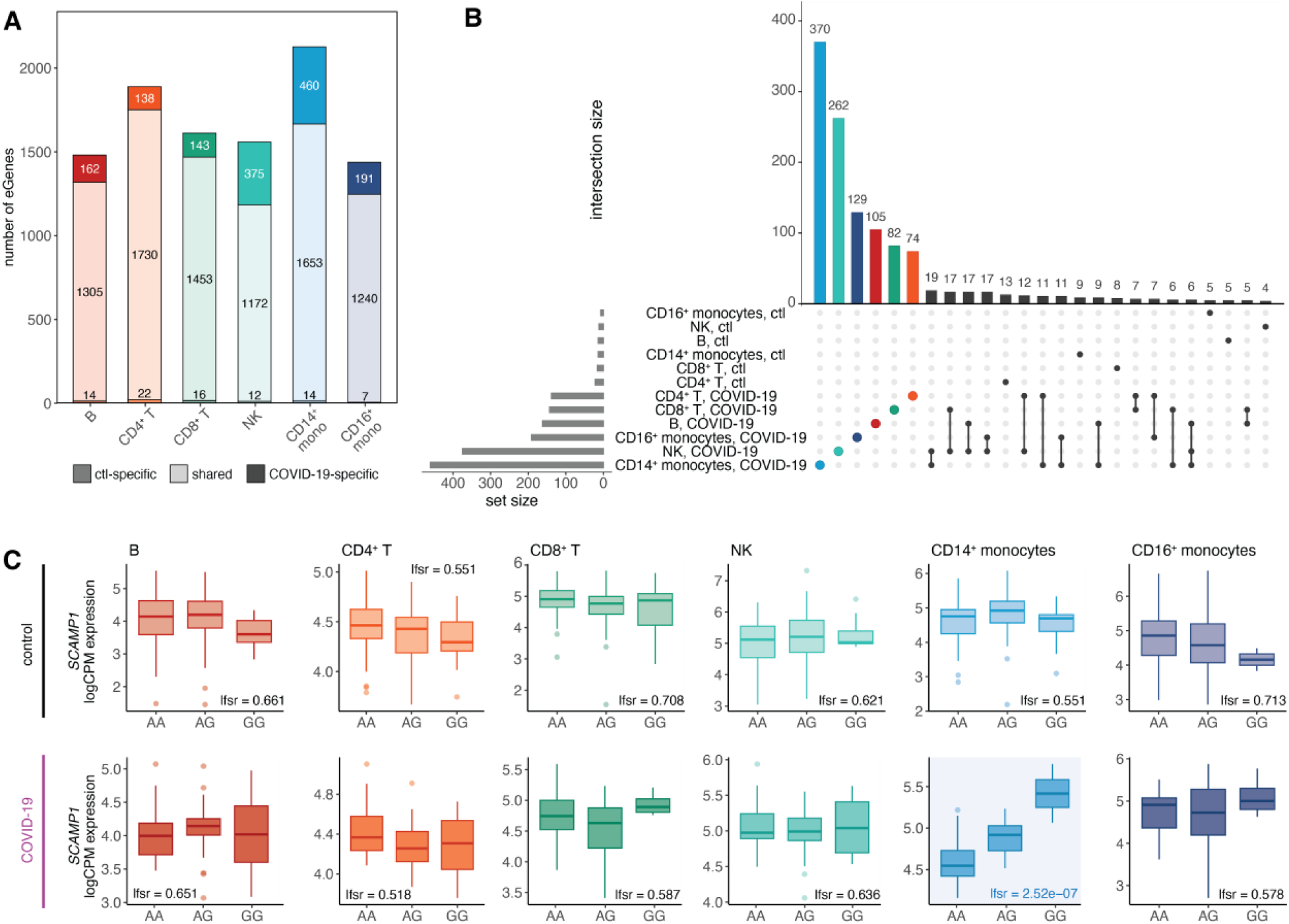
*Cis*-regulatory effects are cell type-specific and disease state-specific. **(A)** Number of shared and disease state-specific eGenes within each cell type. **(B)** Significant condition-specific eGene (lfsr_CTL_ < 0.10 and lfsr_COVID_ > 0.30, lfsr_COVID_ < 0.10 and lfsr_CTL_ > 0.30) sharing patterns across cell types in healthy controls and COVID-19 patients. Patient-specific eGene sets are highlighted by color per cell type. **(C)** Example of a patient-specific genetic effect (i.e., SARS-CoV-2 response eQTL) present only in CD14^+^ monocytes in the gene *SCAMP1* (healthy controls, top plots; COVID-19 patients, bottom plots).

In stark contrast, some cell types, particularly CD14^+^ monocytes and NK cells, displayed a substantial proportion of condition-specific eGenes, where genetic effects were observed exclusively in either control or COVID-19 conditions. Notably, CD14^+^ monocytes and NK cells displayed the greatest fraction of infection-dependent genetic effects (24.8% in NK cells and 22.3% in CD14^+^ monocytes), much higher than the average of 11.0% in other cell types. Strikingly, across all cell types, the overwhelming majority of condition-specific genetic effects (86.3–97.0%) were eQTL observed exclusively in COVID-19 patients rather than in healthy individuals, underscoring the virus’s profound impact on the genetic regulation of immune responses (Fig. 3A).

Condition-specific eGenes were highly cell type-specific, with monocytes possessing a particularly large number of COVID-19-specific eGenes (CD14^+^ monocytes n = 370, CD16^+^ monocytes n = 129), further highlighting the abundance of SARS-CoV-2 response eQTL in the myeloid lineage (Fig. 3B). One prime example of a monocyte-specific response eQTL is the top *cis*-eQTL for *SCAMP1* (rs6453393), a gene involved in cytokine secretion, vesicular trafficking, and membrane transport^26^. This variant exhibited a strong genetic effect unique to CD14^+^ monocytes in COVID-19 patients (lfsr = 2.5 x 10⁻⁷), but no significant effect in other cell types or conditions (lfsr > 0.50) (Fig. 3C). These findings highlight the crucial role of genetic factors in shaping the monocyte response to SARS-CoV-2 infection *in vivo*.

### Response eQTL effects are substantially weaker in the innate immune cells of recovered individuals

Given the abundance of disease state-specific regulatory variation present in COVID-19 patients and absent in healthy individuals, we hypothesized that these eGenes detected only in patients may represent genetic effects only observed during the active infection state. To test whether these genetic effects disappear as infection resolves, we mapped *cis*-eQTL in our cohort of recovered COVID-19 patients who were resampled at various time points following primary SARS-CoV-2 infection (DSO > 20 days, n = 39). We then focused on the innate immune cell compartment (i.e., monocytes and NK cells) to determine how disease state-specific regulatory variation may shift in the convalescent period, as these cell types displayed the greatest number of SARS-CoV-2 response-specific genetic effects (‘response eGenes’, n reQTL: 370 in CD14^+^ monocytes, 262 in NK cells, and 129 in CD16^+^ monocytes) (Fig. 3B). Among these cell type-specific response eGene sets, effect sizes were significantly higher in acute patients compared to follow-ups (p < 1 x 10^-10^ in all three cell types). Indeed, for many eGenes, the effect sizes in follow-up individuals reverted back to the magnitude observed in healthy controls (Fig. 4A), an outcome that was seen across cell types. This result held true even after adjusting for sample size differences across disease state groups and when focusing on the 21 individuals with paired acute and follow-up samples (Fig. S3A).

**Fig. 4.**
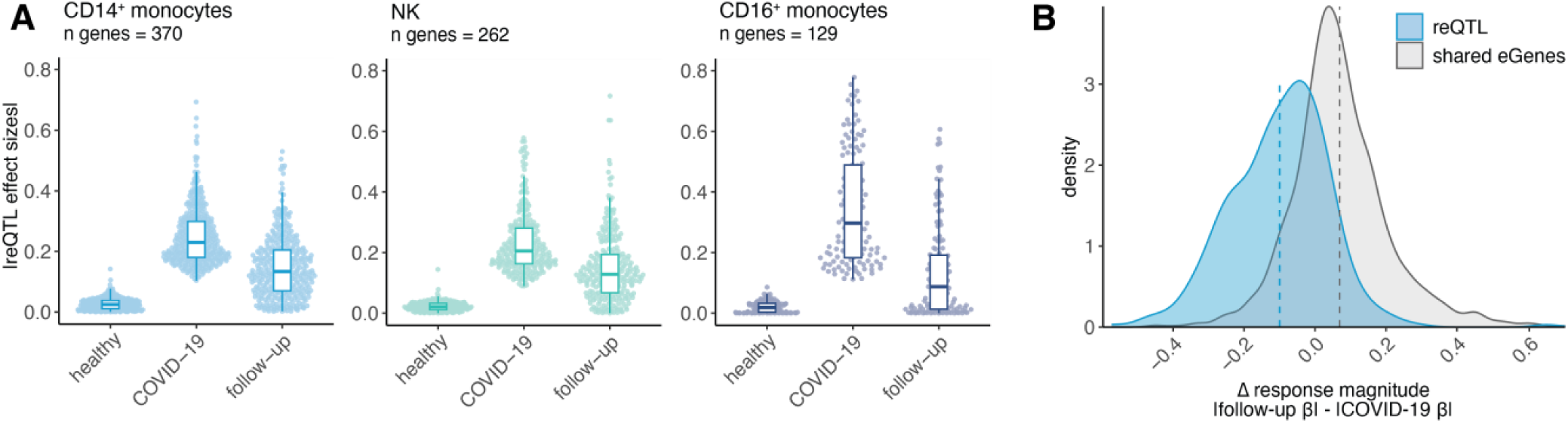
SARS-CoV-2 response eQTL effects revert to baseline in longitudinal follow-up samples. **(A)** Effect sizes for the cell type-specific reQTL gene sets plotted across innate immune cell types in healthy controls, patients, and follow-up samples. All eQTL effect sizes correspond to mash posterior effect sizes. **(B)** Distribution of the change in eQTL effect sizes between follow-up and patient samples (defined as |follow-up β_eQTL_| − |COVID-19 patient β_eQTL_|) for response eQTL (n = 370, blue) and shared eQTL (n = 1,653, gray) in CD14^+^ monocytes. Dashed lines represent the mean Δ response magnitude for the respective gene sets.

To explicitly measure the extent of reQTL effect size reversion coinciding with recovery, we calculated a paired ΔreQTL metric, defined as the difference in magnitude of a response eGene’s effect size in follow-ups compared to COVID-19 patients (i.e., |follow-up β_reQTL_| - |COVID-19 patient β_reQTL_|) specifically in CD14^+^ monocytes. Here, we considered only the effect size magnitude because the vast majority of response eGenes had effect sizes with concordant signs in the patient and follow-up groups (Fig. S3B). For comparison, we also computed this change in response magnitude for the set of shared eGenes between COVID-19 patients and healthy controls (n = 1,653). The mean ΔreQTL for response eGenes was below zero (mean ΔreQTL = −0.10), substantially lower than that for shared eGenes (mean = 0.07) (Fig. 4B). This value was also significantly lower than expected by chance (p < 0.001), as determined by randomly sampling the same number of genes (n = 370) from shared CD14^+^ monocyte eGenes 1,000 times (Fig. S3C). These results indicate that infection mediates dynamic genetic effects and plays a significant role in disease state-dependent gene-environment interactions.

### Cell state-dependent *cis*-regulatory effects are prevalent in CD14^+^ monocytes and can capture clinical features of patient cohorts

We identified several immune and metabolism-related pathways, including TNF-α signaling via NF-κB, oxidative phosphorylation, IFN-γ and IFN-α responses, inflammatory response, and apoptosis, as being strongly associated with disease severity across multiple cell types in COVID-19 patients (Fig. 2D). Given this, we hypothesized that some of the patient-specific genetic effects detected might be driven by heterogeneity in functional cell states within these clinically relevant pathways. To determine whether cell states defined at the single-cell level are dynamically regulated by *cis* variation, we directly mapped single-cell eQTL in COVID-19 patients using our comprehensive single-cell data.

To measure cell state-dependent *cis*-regulatory effects, we applied a continuous measure of cell state, which has been shown to capture more state-dependent regulatory variation than discrete classifications. For each pathway, we calculated a numeric score summarizing the activity for each single cell using ssGSEA (see Methods for details). To map continuous state-dependent *cis*-eQTLs within each cell type, we used a poisson mixed-effects interaction model, a method that has proven successful in identifying state-dependent eQTLs in CD4^+^ T cells^7^. This model tests for genotype-cell state interactions by modeling unique molecular identifier (UMI) counts per gene as a function of genotype at the eQTL variant. We controlled for donor- and cell-level covariates, including age, sex, gene expression PCs, genotype PCs, total UMI count, and mitochondrial UMI percentage (illustrated in Fig. 5A).

**Fig. 5.**
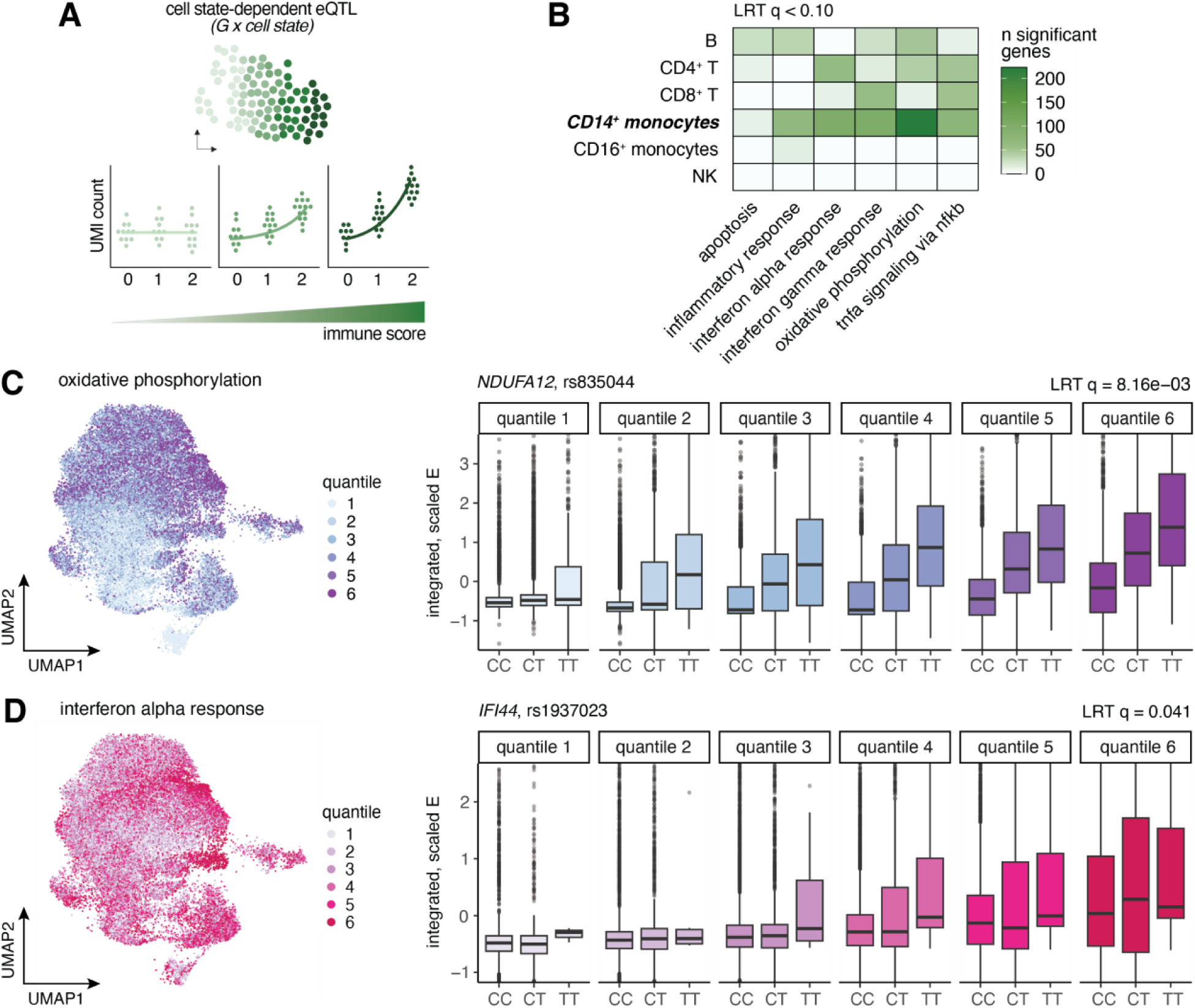
Cell state-dependent single-cell eQTL are prevalent, particularly in CD14^+^ monocytes of COVID-19 patients. **(A)** Schematic of how cell state-dependent single-cell eQTL were evaluated. For each cell type independently, a poisson mixed effects (PME) model was fit to the UMI counts for each gene, correcting for various biological and technical covariates, to test for the interaction between genotype (0, 1, 2) and various functional cell state scores (represented by the green gradient bar). In this instance, no genetic effect is observed among cells with a low immune score (light green), whereas cells with a high immune score display a substantially larger genotype effect (dark green). **(B)** Number of significant cell state-dependent eQTL (LRT q-value < 0.10) for each functional cell state tested (x-axis) across cell types. **(C-D)** UMAP visualizations of all CD14^+^ monocytes in COVID-19 patients colored by **(C)** oxidative phosphorylation score quantiles and **(D)** IFN-α response score quantiles (left), and examples of cell state-dependent eQTL for each of the corresponding functional pathways (right). In these examples, single-cell gene expression estimates (y-axis) are plotted by genotype and binned by cell state score quantiles for each visualization, although we treated cell state as a continuous variable in our models. The quantiles shown directly correspond to the UMAP quantile scale.

For each cell type, we focused on the top gene-SNP pairs identified as eQTLs in COVID-19 patients from the pseudobulk analysis (ranging from 1,395 genes in B cells to 2,084 genes in CD14^+^ monocytes) to assess cell state-dependent genotype effects. Of the six pathways considered, we detected 1,022 significant cell state-dependent interactions with genotype (likelihood ratio test [LRT] q value < 0.10) across all cell type and cell state combinations, mapping to 468 unique eGenes total (17.2% of tested genes) (Fig. 5B, Table S6). CD14^+^ monocytes displayed the largest number of cell state-dependent eQTL across pathways (n = 569 eGenes), while other cell types exhibited more modest state-dependent effects (n = 0 – 171 eGenes). In CD14^+^ monocytes, five of the six pathways were associated with over 50 state-dependent loci, including oxidative phosphorylation (n = 223), IFN-α response (n = 99), IFN-γ response (n = 98), TNF-α signaling via NF-κB (n = 73), and inflammatory response (n = 66) (Fig. 5B).

Oxidative phosphorylation stood out as the functional state most associated with dynamic state-dependent genetic effects, with 223 eGenes detected, corresponding to 10.7% of those tested. One of the top oxidative phosphorylation-dependent variants was rs835044 (LRT q = 8.2 x 10^-3^), a lead *cis*-eQTL 2 kb upstream of *NDUFA12*, a gene encoding the A12 subunit of mitochondrial complex I^27^, which shows a strong genetic effect in cells with high oxidative phosphorylation scores (quantiles 4 - 6) but virtually no genetic effect in cells with low scores (quantile 1) (Fig. 5C). Loss-of-function variants in *NDUFA12* have been linked to a wide array of clinical phenotypes, most frequently a progressive neurodegenerative disorder known as Leigh syndrome^27,28^, suggesting that variation in A12 subunit levels can have substantial clinical consequences.

Many cell state-dependent eQTL were also found for the IFN-α and IFN-γ response pathways, with 4.0% and 3.9% of tested eGenes showing state-dependent genetic variation, respectively. One such variant was rs1937023, a lead *cis*-eQTL upstream of *IFI44*, an interferon-stimulated gene encoding interferon-induced protein 44, which only displays a genetic effect in cells with high IFN-α response scores (LRT q = 0.041) (Fig. 5D). Experimental knockout of *IFI44* in mammalian airway epithelial cells led to increased respiratory syncytial virus (RSV) titers^29^, suggesting that variation in *IFI44* levels can have functional repercussions specifically in the context of viral infection.

### *Cis*-genetic signals colocalize with COVID-19 disease severity risk loci exclusively in COVID-19 patients

Genome-wide association studies (GWAS) provide a means to link regions of the genome with particular traits of interest, giving us the ability to uncover associations with complex disease phenotypes. *Cis*-genetic effects that colocalize with GWAS signals are strongly enriched for causal drivers of variation in disease susceptibility across individuals^30^. To evaluate whether our response eQTL may mechanistically underlie any known COVID-19 GWAS risk loci, we performed colocalization analysis using GWAS results derived from the COVID-19 Host Genetics Initiative^11^, a consortium that has conducted the largest COVID-19 GWAS to date^13^. We integrated our eQTL mapping data in healthy controls, COVID-19 patients, and follow-ups across cell types with two GWAS meta-analyses for COVID-19 disease severity phenotypes: critical illness (A2, very severe respiratory-confirmed COVID-19 versus population) and hospitalization (B2, hospitalized versus population)^11^ to test for common etiological genetic signals.

Across cell types and disease states, we detected 19 signals across 6 unique eGenes that significantly colocalized (posterior probability of colocalization [PP4] > 0.80) with critical illness or hospitalization GWAS risk loci (defined as GWAS SNP meta p-value < 1 x 10^-4^) (Table S7). Of these eGenes, 50% (3 out of 6) colocalized with eQTL exclusively found in COVID-19 patients: *IFNAR2* in CD4^+^ T cells (critical illness and hospitalization GWAS), *JAK1* in CD16^+^ monocytes (hospitalization GWAS only), and *SNRPD2* in CD14^+^ monocytes (hospitalization GWAS only). Notably, two of these genes, *JAK1* and *IFNAR2,* are key canonical mediators of the immune response, both playing critical roles in cytokine signal transduction and interferon response pathways^31,32^. The lead SNP driving the colocalization signal for *IFNAR2* in CD4^+^ T cells of patients, rs9636867 (PP4_A2_ = 0.84, PP4_B2_ = 0.84) (Fig. 6A, right), has previously been shown to colocalize for severe COVID-19 outcomes in whole blood and CD4^+^ T cells of COVID-19 patients and was estimated to be causal^33,34^. This colocalization signature was noticeably absent in healthy controls (Fig. 6A, left) and in follow-ups (Fig. S4A). Similarly, the lead SNP driving the eQTL signal in *SNRPD2*, rs7246757, colocalized in CD14^+^ monocytes of acute COVID-19 patients (PP4_B2_ = 0.87) (Fig. 6B, right), and the gene itself has been implicated as a protein-protein interaction network hub gene associated with SARS-CoV-2 infection^35^. Again, this colocalization signature was entirely absent in control (Fig. 6B, left) and follow-up samples (Fig. S4B), indicating that variation in severe COVID-19 outcomes may, in part, be due to *cis*-regulatory variants that exert their effects in disease-specific and cell type-specific manners.

**Fig. 6.**
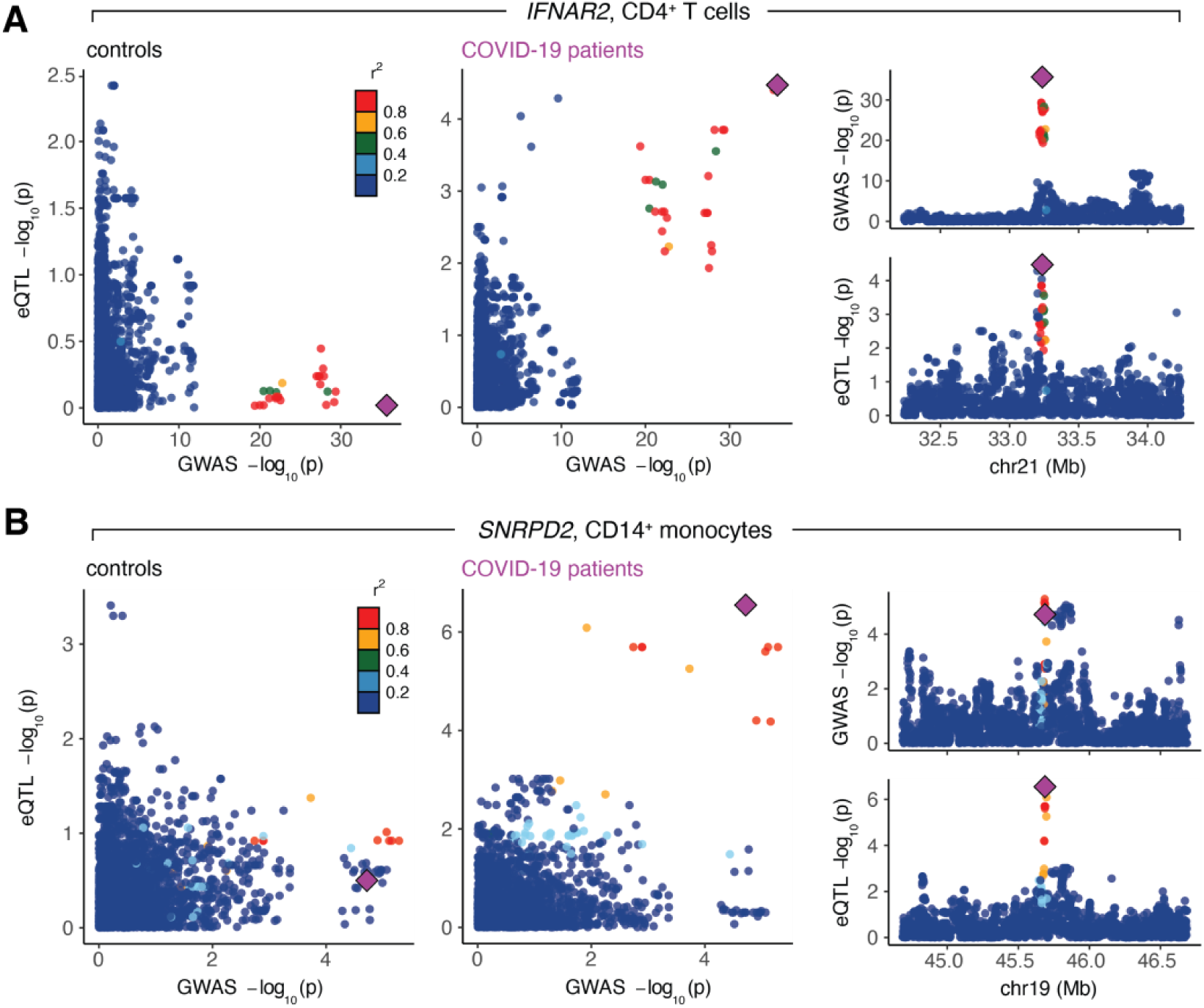
Colocalization signals for COVID-19 disease severity phenotypes are specific to COVID-19 patients. **(A)** The lead SNP for *IFNAR2*, rs9636867, colocalizes in CD4^+^ T cells for hospitalization due to severe COVID-19 in patients (right) but not controls (left). **(B)** The lead SNP for *SNRPD2*, rs7246757, colocalizes in CD14^+^ monocytes for hospitalization due to severe COVID-19 in patients (right) but not controls (left). For both **(A)** and **(B)**, the larger plots on the left show the correlation between GWAS p-values (x-axis) and eQTL p-values (y-axis) in controls and patients. The smaller plots on the right show Manhattan plots for the GWAS signal (top) and the eQTL signal in the COVID-19 patients (bottom). The lead SNP is depicted as a purple diamond.

## Discussion

Prior studies have leveraged *in vitro* pathogen challenges and immune stimulations to probe gene regulatory variation in cells, reporting hundreds of response eQTL in different infection contexts^1–5,36,37^. This experimental approach involves the isolation and culture of primary immune cells from healthy donors, which are then subsequently challenged in laboratory settings. Unlike previous immune response eQTL studies, here we measure genetic effects directly in cells derived from patients responding to a pathogen, revealing considerable context specificity in genetic regulation that is arguably more relevant to disease associations than that measured in controlled *in vitro* systems. We show that cell type-specific, disease state-specific, and cell state-dependent genetic variation is abundant, affecting 25.6% of all genes tested across cell types and disease states and is particularly common in CD14^+^ monocytes and NK cells. Further, we establish that single cells can harbor distinct genetic effects that are dependent on their underlying immunological or metabolic functional states and that, in certain cases, these continuous states are associated with clinical features of patients. More broadly, genetic interaction effects likely play a role in dynamically modulating immune responses throughout the course of an infection and may also contribute to differential disease outcomes, especially considering the fact that monocytes, and more generally cells in the myeloid compartment, are susceptible to immune dysregulation following SARS-CoV-2 infection^38,39^.

Of particular clinical interest is biological variation in the interferon response, a critical antiviral pathway induced upon the detection of viral pattern recognition receptors. This response involves the induction of IFNs, a group of cytokines that directly inhibit viral replication and activate bystander immune cells, such as dendritic cells and monocytes^40^. Variation in the timing and magnitude of the IFN response across individuals is well-documented, particularly in the context of SARS-CoV-2 infection^41–44^. Multiple studies have linked this variation with differences in COVID-19 severity and disease progression, revealing a dual role for IFNs in the clinical course of COVID-19^45,46^. In the blood, the upregulation of type I IFNs and IFN-stimulated genes (ISGs) shortly after initial infection is associated with protection^21^, but their delayed induction is a hallmark of severe disease^47–49^. Sustained IFN signaling has also been shown to inhibit the development of appropriate antibody responses, ultimately leading to increased disease pathology and severity^44^. We also observe a relationship between IFN signaling and severity, with milder COVID-19 cases displaying elevated expression of IFN-α and IFN-γ response genes specifically in T cells and CD14^+^ monocytes.

Of note, we detect 224 eGenes (47.8% of all state-dependent eGenes identified) across cell types with expression levels simultaneously dependent on both underlying genetic variation and the magnitude of the IFN response itself, revealing it to be one of the pathways most associated with cell state-dependent genetic interaction effects. This finding only adds to the complexity of how dynamic immune response variation is connected to variation in molecular traits, here through an interaction with host genetics, which may ultimately have downstream effects on disease phenotypes. Indeed, we find that IFN response scores calculated at the single-cell level correlate with patient severity. Together, our results argue that gene-environment interactions are abundant and likely play a direct role in the clinical setting.

While we identify only a handful of colocalizing eQTL, of the eGenes that colocalize with COVID-19 disease severity phenotypes, half are detected only in COVID-19 patients, indicating that SARS-CoV-2 infection is necessary to induce these signals. Similar disease state-dependent colocalization has been described previously, with the variant rs8176719 colocalizing only in T effector memory cells 16 hours post-anti-CD3/CD28 stimulation for both severity and susceptibility COVID-19 GWAS at the *RALGDS2* locus^33^. In the same study, the intronic risk variant in *IFNAR2*, rs9636867, the same lead SNP-eGene pair for which we identify a patient-specific colocalization signal in CD4^+^ T cells, colocalized with severe COVID-19 disease only for symptomatic individuals who were SARS-CoV-2^+^ in CD4^+^ T cells^33^. A different COVID-19-associated intronic risk variant in *IFNAR2*, rs13050728, has also been shown to increase *IFNAR2* expression in classical monocytes specifically in COVID-19 patients compared to healthy controls in an independent study^14^.

These findings highlight the role that context specificity plays in the genetic regulation of disease associated-traits and stress the importance of measuring molecular phenotypes in pertinent environmental conditions and cell types. They also raise the question of how gene-environment interactions may contribute to the problem of missing heritability, the phenomenon in which only a small fraction of overall trait heritability is explained by trait-associated variants^50,51^. Although trait-associated loci are enriched for eQTL^52^, only ∼40% of GWAS variants colocalize with eQTL in relevant tissues, which drops to ∼20% for autoimmune trait GWAS^53,54^. More recently, trait mapping studies have been extended to incorporate a larger array of quantitative traits, including alternative splicing^55^, chromatin accessibility, and histone modification levels^37^. The inclusion of alternative regulatory mechanisms has significantly increased the number of colocalizing loci and heritability estimates of GWAS phenotypes, yet a large proportion of heritability remains unexplained, potentially due to context-specific gene-environment interactions.

Although we have described how gene-environment interactions can shape immune responses in one specific viral infection setting, it is necessary to define how such effects contribute to a wider range of disease states and environmental contexts to better understand the genetic and environmental underpinnings of immune response variation across individuals. As the number of patient cohorts with single-cell phenotyping and genotyping data rise, it will be important to extend this framework to other single-cell eQTL mapping studies to measure the full extent of cell state-dependent regulatory heterogeneity.

## Acknowledgements

We thank all study participants and the clinical research teams for their contributions. We thank members of the Barreiro lab for their constructive comments and feedback. This work was completed in part with resources provided by the University of Chicago Research Computing Center. Support from Calcul Québec and Compute Canada is additionally acknowledged. We thank the University of Chicago Genomics Facility (RRID: SCR_019196) for their assistance with sequencing. Figures 1A and 5A were created with BioRender.com.

## Author contributions

L.B.B. supervised the study. H.E.R. and L.B.B. designed the experiments. E.B.R., T.N., C.L., A.P., N.A., A.F., M.D., J.B.R., and D.E.K. provided patient samples and curated data. H.E.R., V.L., E.K., C.B., and A.D. performed the experiments and sample collections. H.E.R., R.A.G., and T.N. performed computational analyses. H.E.R. and L.B.B. wrote the manuscript, with input from all authors.

## Data and materials availability

Further information and requests for resources should be directed to Luis B. Barreiro. Raw data contained in BQC19, including whole genome sequencing files, are stored on SecureData4Health (https://www.sd4health.ca/), and are accessible via BQC19’s access procedures. To access these files, a data access request must be submitted. Instructions on how to submit this request are available at https://www.bqc19.ca/en/access-data. Researchers from both academia and private entities are eligible to apply, and the research project must be approved by a research ethics board. A Data Access Committee will then review the application, and the data will be made available to the applicant upon approval.

## Supplementary materials

Materials and Methods

Figures S1-S4

Tables S1-S8

## Materials and Methods

### Participants and samples

We prospectively investigated hospitalized COVID-19 patients between April 2020 and December 2021 who initially presented with a symptomatic infection and positive SARS-CoV-2 nasopharyngeal swab polymerase chain reaction. All participants were admitted to the Centre Hospitalier de l’Université de Montréal (CHUM) and recruited into the Biobanque Québécoise de la COVID-19 (BQC19)^56^. Patients had no known prior exposure to SARS-CoV-2 (i.e., all infections were primary infections), were not vaccinated at the time of primary sampling (days after symptom onset [DSO] ≤ 20), and did not undergo plasma transfer therapy. Blood draws were performed during the acute phase of SARS-CoV-2 infection (defined as DSO ≤ 20 days, mean DSO = 12.1 days, DSO range = 6 - 20 days, n = 63 samples) and during various convalescent follow-up time points (defined as DSO > 20 days, mean DSO = 128.8 days, DSO range = 31 - 370 days) for a subset of individuals sampled during the acute phase (n = 39 samples). Additionally, PBMCs collected prior to the COVID-19 pandemic from healthy control individuals living in Montréal, Canada (n = 18 samples) were processed for single-cell data collection in parallel with infected patient samples. We also computationally integrated a set of publicly available healthy controls (n = 90 individuals) described in Randolph et al. (2021)^4^, which is detailed below (“Single-cell RNA-sequencing data processing and integration”). The study was approved by the respective IRBs (multicentric protocol: MP-02-2020-8929 for BQC19 participants; CHUM protocol 19.387 for control individuals) and written, informed consent was obtained from all participants or, when incapacitated, their legal guardian before enrollment and sample collection.

### DNA sequencing and imputation

DNA was extracted from whole blood using the Chemagic™ DNA Blood 400 H96 kit (Perkin Elmer, CMG-1091). SNP genotyping was conducted using the Axiom™ Precision Medicine Research Array from Applied Biosystems (Applied Biosystems, 902981) per the manufacturer’s instructions. The array was processed using the GeneTitan™ Multi-Channel instrument (Applied Biosystems). All samples were grouped with the Axiom Analysis Suite 5.1.1 software, and the “Best Practice Workflow” was performed using the following high-quality call rate parameters: Axiom_PMRA.r3 library and threshold configuration Human.v5 with minimum call rate of 97.0%. Marker quality control tests were performed on a subset of ancestrally homogeneous participants, who were determined via comparison to 2,504 individuals across 5 super populations from the 1000 Genomes Project Phase 3 data ^57^. Batch effect quality control and replicate discordance checks were performed, and variants that failed either test were removed. Only single nucleotide variants with single character allele-codes (A, C, G, or T) (PLINK --snps-only ‘just-acgt’ option) were retained. Additionally, variants with low allele frequencies (minor allele frequency [MAF] < 0.001), low genotyping call rates (marker-wise missingness < 0.01), a deviation from Hardy-Weinberg equilibrium (HWE) (p-value < 1×10^-6^), and positioned in regions of high link disequilibrium (LD) were removed.

Sample quality filtering was performed considering the set of filtered genotypes described above. Outlier samples with a high genotype missingness rate (overall missing genotype rate > 0.04) or high/low principal component corrected heterozygosity rate on autosomal chromosomes (> ±3SD, respectively) were considered low quality and removed. Sex chromosome composition was determined by estimating X chromosome marker heterozygosity using PLINK (--check-sex 0.4 0.7). Individuals with discordant self-reported sex and genetic sex were removed prior to genotype imputation. All other samples that passed quality control filters were used for imputation. Genotype phasing and imputation was performed using the Michigan Imputation Server^58^ with the TOPMed reference panel^59^. After imputation, variants with a posterior genotype probability (GP) 90% were set to missing within each individual using QCTOOL (v2.0.7, -threshold 0.9 filter).

### Whole blood processing

At the time of sampling, whole blood was collected in up to three tubes containing acid citrate dextrose (ACD) and processed within 6 hours of collection. Blood from the same donor was pooled and centrifuged at 400 g for 10 min at room temperature (RT). After centrifugation, plasma was collected, aliquoted, and stored at ™80°C. The remaining blood was topped up to 30 ml with HBSS medium at RT. Ficoll-Paque separation was then used to isolate PBMCs. PBMCs were washed with R+ (RPMI 1640 + 0.1M HEPES + 20 U/ml Penicillin-Streptomycin), resuspended in 5 ml R+ with 10% fetal bovine serum (FBS), and counted with Trypan blue. Cells were spun down at 400 g for 10 min at 4°C and resuspended in cold FBS at 20 M/ml. A freezing solution of FBS with 20% DMSO was added drop-by-drop to the cell suspension while the tube was continuously agitated. Cell suspensions were transferred into cryovials (1 ml/vial), immediately placed into Mr. Frosty Freezing Containers, and stored at −80°C. The following day, PBMCs were transferred to liquid nitrogen for long-term storage.

### Sample processing for single-cell RNA-sequencing

PBMCs were thawed in groups of 3 to 4 samples (processing batch 1) or 16 to 19 samples (processing batch 2), rested for 2 hours in RPMI 1640 supplemented with 10% FBS (Corning, MT35015CV), 2 mM L-glutamine (ThermoFisher Scientific, 25-030-081), and 10 ug/ml gentamicin (ThermoFisher Scientific, 15710064), and subsequently processed for single-cell collection. Cells from different samples were pooled per processing batch for a total of 29 multiplexed sample batches (n = 124 samples). For each multiplexed cell pool, 12,000 cells were targeted for collection using the Chromium Next GEM Single Cell 3’ Reagent (v3.1 Dual Index chemistry) kit (10x Genomics, 1000268). After GEM generation, the reverse transcription (RT) reaction was performed in a thermal cycler as described (53°C for 45 min, 85°C for 5 min), and post-RT products were stored at −20°C for up to one week until downstream processing.

### Single-cell RNA-sequencing library preparation and sequencing

Post-RT reaction cleanup, cDNA amplification and sequencing library preparation were performed as described in the Single Cell 3’ Reagent Kits v3.1 (Dual Index) User Guide (10x Genomics). Briefly, cDNA was cleaned with DynaBeads MyOne SILANE beads (ThermoFisher Scientific, 37002D) and amplified in a thermal cycler using the following program: 98°C for 3 min, [98°C for 15 s, 63°C for 20 s, 72°C for 1 min] x 11 cycles, 72°C 1 min. After cleanup with the SPRIselect reagent kit (Beckman Coulter, B23317), libraries were constructed by performing the following steps: fragmentation, end-repair, A-tailing, double-sided SPRIselect cleanup, adaptor ligation, SPRIselect cleanup, sample index PCR (98°C for 45 s, [98°C for 20 s, 54°C for 30 s, 72°C for 20 s] x 14 cycles, 72°C 1 min), and double-sided SPRIselect size selection. Prior to sequencing, all multiplexed single-cell libraries were quantified using the KAPA Library Quantification Kit for Illumina Platforms (Roche, 50-196-5234). For each processing batch (n = 2), libraries were pooled in an equimolar ratio and sequenced 100 base pair paired-end on an Illumina NovaSeq 6000 (processing batch 1 average mean reads per cell = 48,613, average median genes detected per cell = 1,627; processing batch 2 average mean reads per cell = 59,246, average median genes detected per cell = 2,007).

### Single-cell RNA-sequencing data processing and integration

FASTQ files from each multiplexed capture library were mapped to the pre-built GRCh38 human reference transcriptome (downloaded 10x Genomics) using the cellranger (v6.0.2) count function^60^. souporcell (v2.0, Singularity v3.4.0)^61^ in --skip_remap mode was used to demultiplex cells into samples based on genotypes from a common variants file (1000 Genomes Project samples filtered to SNPs with ≥ 2% allele frequency in the population, downloaded from https://github.com/wheaton5/souporcell). For each sample batch, hierarchical clustering of the known genotypes obtained from DNA-sequencing and cluster genotypes estimated by souporcell was used to assign individuals to souporcell cell clusters. All samples except for three were successfully demultiplexed; samples unable to be confidently assigned to a set of cells were removed (n samples retained = 121). After demultiplexing, Seurat (v4.3.0, R v4.0.3)^62^ was used to perform cell-level quality control filtering. One sample was removed due to a very low number of cells captured (n = 20 cells total), leaving a total of 120 samples. High-quality cells were retained for downstream analysis if they had: 1) a “singlet” status called by souporcell, 2) between 500 – 4000 genes detected (nFeature_RNA), 3) a mitochondrial UMI percentage < 20%, and 4) less than 25,000 total molecules (nCount_RNA), leaving 236,143 cells. Gene filtering was performed using the CreateSeuratObject min.cells parameter, in which only genes present in at least five cells were kept (n = 30,986 genes).

Due to the large discrepancy between the number of cells assayed in healthy control individuals (n = 38,663) versus acute and convalescent samples (n = 197,480) in our dataset, we integrated a publicly available set of high-quality cells derived from control, non-infected individuals (n = 124,976 cells, 90 samples) described in Randolph et al., (2021)^4^, hereafter referred to as the “non-infected IAV controls”. First, we removed IAV-derived transcripts (n = 10 genes) from the raw count matrix of the non-infected IAV controls. Next, we merged all datasets, split the resulting Seurat object by dataset (“COVID batch1”, “COVID batch2” or “IAV controls”), and ran SCTransform^63^ to normalize and scale the UMI counts within dataset. We simultaneously regressed out variables corresponding to experiment batch, percent mitochondrial UMIs per cell, and individual label in all datasets, and additionally, regressed out sampling time point (e.g., control, acute, follow-up) in the COVID data. We then integrated the three datasets together using the SelectIntegrationFeatures, PrepSCTIntegration, FindIntegrationAnchors, and IntegrateData framework^62^. After integration, dimensionality reduction was performed via UMAP (RunUMAP function, dims = 1:30) and PCA (RunPCA function, npcs = 30). A Shared Nearest Neighbor Graph was constructed using the FindNeighbors function (dims = 1:20, all other parameters set to default), and clusters were subsequently called using the FindClusters algorithm (resolution = 0.5, all other parameters set to default)^62^. In total, our integrated dataset consisted of 361,119 high-quality cells across all samples (n = 236,143 from the combined COVID datasets, n = 124,976 from the non-infected IAV dataset, n = 208 samples altogether).

### Cell type assignment

We performed cell type annotation via label transfer to map cell type information onto our data. To perform the label transfer, we downloaded a multimodal human PBMC reference dataset derived from scRNA-seq paired with CITE-seq as described in Hao et al.^15^. We followed the Seurat v4 Reference Mapping workflow, consisting of the FindTransferAnchors and MapQuery functions, with the Hao et al. reference dataset used as our reference UMAP and the following parameters: normalization.method = “SCT” and reference.reduction = “spca”. These fine-scale populations were then collapsed into the following broad super populations encompassing the six major cell types found in PBMCs using the predicted.celltype.l2 definitions derived from Hao et al.: CD4^+^ T cells = c(“CD4 CTL”, “CD4 Naive”, “CD4 Proliferating”, “CD4 TCM”, “CD4 TEM”, “Treg”), CD8^+^ T cells = c(“CD8 Naive”, “CD8 Proliferating”, “CD8 TCM”, “CD8 TEM”), NK cells = c(“NK”, “NK Proliferating”, “NK_CD56bright”), CD14^+^ monocytes = “CD14_monocytes”, CD16^+^ monocytes = “CD16_monocytes”, and B cells = c(“B intermediate”, “B memory”, “B naive”). In total, we annotated 342,127 high-quality cells falling into the major PBMC populations across all individuals and conditions (n CD4^+^ T cells = 153,479, CD8^+^ T cells = 53,562, CD14^+^ monocytes = 70,060, CD16^+^ monocytes = 5,446, B cells = 34,805, NK cells = 24,775).

### Calculation of pseudobulk estimates

Pseudobulk estimates were used to summarize single-cell expression values into bulk-like expression estimates within samples. This was performed for all six major cell types (CD4^+^ T cells, CD8^+^ T cells, B cells, CD14^+^ monocytes, CD16^+^ monocytes, NK cells). Within each cell type cluster for each sample, raw UMI counts were summed across all cells assigned to that sample for each gene using the sparse_Sums function in textTinyR (v1.1.3) (https://cran.r-project.org/web/packages/textTinyR/textTinyR.pdf), yielding an n x m expression matrix, where n is the number of samples included in the study (n = 208) and m is the number of genes detected in the single-cell analysis (m = 30,986) for each of the 6 clusters.

### Calculation of residuals for modelin

For each cell type, lowly-expressed genes were filtered using cell type-specific cutoffs (removed if they had a median logCPM < 1.0 in CD14^+^ monocytes, < 1.5 in CD4^+^ T cells, < 2.0 in B cells and CD8^+^ T cells, < 2.5 in CD16^+^ monocytes, and < 3.0 in NK cells), leaving the following number of genes per cell type: CD4^+^ T cells = 10,337, CD8^+^ T cells = 10,036, B cells = 10,179, CD14^+^ monocytes = 10,882, CD16^+^ monocytes = 9,398, and NK cells = 9,882. Within each cell type, only samples with ≥ 5 cells per sample were kept for downstream modeling. Further, three samples were removed for downstream analysis because they consistently clustered as outliers on gene expression PCAs for multiple cell types (one COVID-19 patient at the acute infection time point and two non-infected IAV controls), leaving the following number of samples per cell type:

| Cell type | N healthy controls | N patients | N follow-ups |
| --- | --- | --- | --- |
| B | 106 | 63 | 38 |
| CD4<sup>+</sup> T | 106 | 63 | 39 |
| CD8<sup>+</sup> T | 106 | 63 | 39 |
| CD14<sup>+</sup> monocytes | 106 | 63 | 39 |
| CD16<sup>+</sup> monocytes | 47 | 44 | 39 |
| NK | 63 | 63 | 39 |

After removing lowly-expressed genes, normalization factors to scale the raw library sizes were calculated using calcNormFactors in edgeR (v 3.26.8)^64^. The voom function in limma (v3.40.6)^65^ was used to apply these size factors, estimate the mean-variance relationship, and convert raw pseudocounts to logCPM values. The inverse variance weights calculated by voom were obtained and included in the respective lmFit call for all downstream models unless otherwise noted^65^.

### Calculation of per-individual ssGSEA scores

To construct the ssGSEA Hallmark pathway scores, we calculated single sample Gene Set Enrichment Analysis (ssGSEA) scores from the pseudobulk COVID-19 patient logCPM gene expression estimates corrected for age, sex, dataset, and the number of cells for a given cell type collected per sample using the Gene Set Variation Analysis (GSVA, v1.32.0) package in R with default parameters and method = “ssgsea”^66^. ssGSEA is a method that allows you to summarize gene expression patterns for any desired target gene set, and for each sample, it will return a score representative of that gene set. These scores were calculated per cell type, and for each of the pathway-specific ssGSEA scores, the input gene set was derived from either a Hallmark or Gene Ontology (GO) Biological Process gene set^22^. The following gene sets were used to define the per-sample pathway scores: (1) inflammatory response score – Hallmark inflammatory response pathway, (2) TNF-α score – Hallmark TNF-α signaling via NF-κB pathway, (3) oxidative phosphorylation score – Hallmark Oxidative phosphorylation pathway, and (4) antigen processing score – GO Biological Process antigen processing and presentation pathway.

### Modeling SARS-CoV-2 infection effects

Only healthy controls and COVID-19 patients sampled during the primary infection time point were retained for modeling of infection effects (i.e., follow-up samples were excluded). The following linear model was used to identify genes differentially expressed between healthy control individuals and COVID-19 patients:

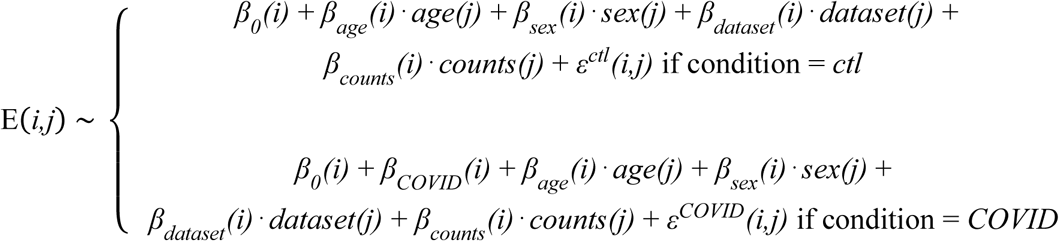

Here, *E(i,j)* represents the expression estimate of gene *i* for individual *j*, *β_0_(i)* is the global intercept accounting for the expected expression of gene *i* in a non-infected female measured in the COVID batch 1 dataset, and *β_COVID_(i)* represents the global estimate of the effect of SARS-CoV-2 infection in patients per gene. Age represents the mean-centered, scaled (mean = 0, sd = 1) age per individual, with *β_age_(i)* being the effect of age on expression levels, sex represents the self-identified sex for each individual (factor levels = “Female”, “Male”), with *β_sex_(i)* capturing the effect of sex on expression, dataset represents the dataset in which the sample was obtained (factor levels = “COVID batch 1”, “COVID batch 2”, “IAV controls”), with *β_dataset_(i)* capturing the dataset effect, and counts represents the number of cells captured within that cell type for sample *j*, with *β_counts_(i)* capturing the effect of cell number on expression. Finally, *ε^cdt^* represents the residuals for each respective condition (control or COVID) for each gene *i*, individual *j* pair. The model was fit using the lmFit and eBayes functions in limma^65^, and the estimates of the global infection effect *β_COVID_(i)* (i.e., the differential expression effects due to SARS-CoV-2 infection) were extracted across all genes along with their corresponding p-values. We controlled for false discovery rates (FDR) using an approach analogous to that of Storey and Tibshirani^2,67^, which derives the distribution of the null model empirically. To obtain a null, we performed 10 permutations, where infection status label (i.e., control/COVID) was permuted across individuals. We considered genes significantly differentially expressed upon infection if they had *β_COVID_* |log_2_FC| > 0.5 and an FDR < 0.05.

### Modeling COVID-19 disease severity effects within patients

To model the effect of COVID-19 disease severity on gene expression, we restricted our analyses to COVID-19 patients sampled during the primary infection time point for which we had information about disease severity (n = 63). Disease severity was assessed using a five-point scale of respiratory support needed at the time of patient sampling that includes the following categories: 0-Moderate = no supplemental oxygen (n = 16); 1-Severe = nasal cannula (n = 17); 2-Critical = non-invasive ventilation (n = 9); 3-Critical = intubation (n = 20); 4-Critical = extracorporeal membrane oxygenation (ECMO) (n = 1). The following model was used to evaluate the effect of severity at the time of patient sampling on expression:

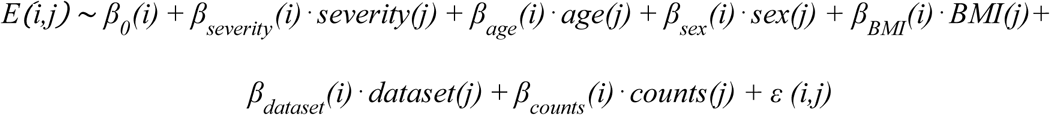

Here, *E(i,j)* represents the expression estimate of gene *i* for individual *j*, *β_0_(i)* is the global intercept accounting for the expected expression of gene *i* in a female COVID-19 patient, and *β_severity_(i)* indicates the effect of severity on gene *i* during the primary sampling time point. Severity (*severity(j)*) represents respiratory support score per individual and was treated as a numeric variable. Body mass index (BMI) represents the mean-centered, scaled (mean = 0, sd = 1) BMI per individual, with *β_BMI_(i)* being the effect of BMI on expression levels. If BMI was not reported for an individual (n missing = 26), this missing data was filled with the average BMI across patients. All other terms in the model are equivalent to that described in “Modeling SARS-CoV-2 infection effects”. The model was fit using the lmFit and eBayes functions in limma^65^, and the estimates of *β_severity_(i)* were extracted across all genes along with their corresponding p-values. We again controlled for false discovery rates (FDR) by empirically deriving the null distribution. To obtain a null, we performed 10 permutations, where respiratory support score (i.e., 0 - 5) was permuted across patients. We considered genes significantly correlated with disease severity if they had an FDR < 0.05.

### Gene set enrichment analyses

The R package fgsea (v1.10.1)^68^ was used to perform gene set enrichment analysis for the severity effects using the H hallmark gene sets^23^. Ranked t-statistics for each cell type were obtained directly from the topTable function in limma^65^, and the background set for a cell type was the set of genes sufficiently expressed (i.e., passed the lowly-expressed gene filter threshold) for that cell type. Pre-ranked t-statistics were used to perform the enrichment using fgsea with the following parameters: minSize = 15, maxSize = 500, nperm = 100,000. Normalized enrichments scores (NES) and Benjamini-Hochberg adjusted p-values output by fgsea were collected for each analysis.

### eQTL mapping and integration with mashr

eQTL mapping was performed for each cell type using the pseudobulk expression data. A linear regression model was used to ascertain associations between SNP genotypes and expression levels. Input expression matrices were quantile-normalized within each set of disease state samples (i.e., healthy controls, acute COVID-19 patients, and follow-ups) prior to association testing. eQTL were mapped separately for each disease state using the R package MatrixEQTL (v2.3)^69^. Prior to mapping, SNPs were filtered using the following criteria in our COVID-19 dataset and the Randolph et al. dataset separately: 1) keep those with a minor allele frequency > 5% across all individuals, 2) exclude those with > 10% of missing data, and 3) exclude those that deviate from Hardy-Weinberg equilibrium at p < 10^-5^ (--maf 0.05 --geno 0.10 --hwe 0.00001 PLINK v1.9 filters)^70^. Only SNPs that passed these filters and were present in both datasets were retained and merged across datasets (n = 4,194,100 SNPs kept). Local associations (i.e., putative *cis*-eQTL) were tested against all SNPs located within the gene body and 100 kilobases upstream and downstream of the transcription start site (TSS) and transcription end site (TES) for each gene tested.

Within our follow-up samples, some individuals were sampled multiple times during the convalescent period. To avoid counting these genetically duplicate samples more than once when eQTL mapping, we downsampled the follow-ups to include only a single sample with DSO > 20 per individual. For each individual with multiple follow-up time points, we chose to keep the sample with the maximum DSO, which dropped our sample size from n = 39 to n = 26. This duplicate sampling structure was not present in the healthy control or acute COVID-19 samples, so the full sample set was used to map eQTL for these disease states.

We accounted for unmeasured surrogate confounders by performing PCA on a correlation matrix based on the gene expression data. Subsequently, up to 15 principal components (PCs) were regressed out prior to performing the association analysis for each gene. A specific number of PCs to regress in each cell type-disease state pair, corresponding to the number of PCs that led to the detection of the largest number of eQTL in each condition, was then chosen empirically (Table S8). To avoid spurious associations resulting from population structure, the first two eigenvectors obtained from a PCA on the genotype data using SNPRelate (v1.20.1, gdsfmt v1.22.0)^71^ were included in the linear model. Other covariates included were age (mean-centered, scaled), sex, number of cells detected per sample, and dataset.

To gain power to detect *cis*-eQTL effects, we implemented mashr^25^, which leverages sharing information across cell types and disease states. We considered a set of shared genes that were expressed across all cell types (n = 7,646). For each of these genes, we chose the single top *cis*-SNP, defined as the SNP with the lowest FDR across all cell types (n = 6) in the acute COVID-19 patient condition, to input into mashr. We extracted the effect sizes and computed the standard errors of these betas from the Matrix eQTL outputs for each gene-SNP pair across cell types and conditions. We defined a set of strong tests (i.e., the 7,646 top gene-SNP associations) as well as a set of random tests, which we obtained from randomly sampling 200,000 rows of a matrix containing all gene-SNP pairs tested merged across conditions. The mashr workflow was as follows: i) the correlation structure among the null tests was learned using the random test subset, ii) the data-driven covariance matrices were learned using the strong test subset (from 5 PCs), iii) the mash model was fit to the random test subset using canonical and data-driven covariance matrices, and iv) the posterior summaries were computed for the strong test subset. We used the local false sign rate (lfsr) to assess significance of our posterior eQTL effects and considered a gene-SNP pair to have a significant eQTL effect if the lfsr was < 0.10.

### Calculation of functional cell state scores per cell

To obtain the cell state scores used for modeling cell state-dependent single-cell eQTL, first, the raw single-cell UMI counts across all samples were obtained per cell type. All subsequent processing steps were performed for each cell type independently. Raw cell counts in the form of a Seurat object were split by dataset, and SCTransform was used to normalize and scale the UMI counts within dataset, regressing the effects of experiment batch, percent mitochondrial UMIs per cell, and age in all datasets, and additionally, sex in the COVID batch 1 and batch 2 datasets. The SelectIntegrationFeatures, PrepSCTIntegration, FindIntegrationAnchors, and IntegrateData pipeline was then used to integrate cells, returning all features following integration (features.to.integrate = all_features)^60^. The scaled data matrix (@scale.data slot) of the integrated data, which holds the residuals of the corrected log-normalized integrated counts, was obtained, and these values were used to calculate ssGSEA scores (using the same parameters described above in “Calculation of per-individual ssGSEA scores”) per cell for our pathways of interest. Here, we applied ssGSEA to the full scaled SCTransform gene *x* cell matrix, allowing us to generate cell state scores for each single cell in the dataset. Our pathways of interest included the following immune-related and metabolism-related pathways in the MSigDB Hallmark gene sets (n = 6)^22^: Apoptosis, Inflammatory response, Interferon-α response, Interferon-γ response, Oxidative phosphorylation, and TNF-α signaling via NF-κB.

### Modeling cell state-genotype interaction effects

We used a poisson mixed effects model to test for cell state-dependent eQTL because this model has previously been used to detect significant cell state-genotype interaction effects in single-cell data^7^. Only COVID-19 patients sampled during the primary infection time point were included in these analyses (n = 63). Single-cell eQTL modeling was performed independently in each cell type; for each cell type, we tested the gene-SNP pairs for which we had evidence of a significant eQTL (lfsr < 0.10) within patients in the pseudobulk eQTL analysis (n genes: B cells = 1,395, CD4^+^ T cells = 1,804, CD8^+^ T cells = 1,508, CD14^+^ monocytes = 2,084, CD16^+^ monocytes = 1,410, NK cells = 1,523). For CD4^+^ T cells, we downsampled the number of cells prior to constructing the model inputs to 60,000 cells due to vector size constraints in R. To control for genetic background and latent confounders, we included both genotype and expression PCs in our cell state eQTL models. We computed genotype PCs using the same approach as above in “eQTL mapping and integration with mashr”. Expression PCs were calculated from non-batch corrected integrated and scaled counts using the same method as described in “Calculation of functional state scores per cell,” but omitting the batch correction step (i.e., no variables were regressed in the SCTransform call). PCA was run on the cell *x* gene matrix of non-corrected integrated and scaled counts subset on the top 3,000 variable features using the prcomp_irlba function in the R package irlba (v2.3.5.1)^72^.

To test for interactions with cell state, we used the following poisson mixed effects interaction model, where each gene’s UMI counts were modeled as a function of genotype as well as additional donor-level and cell-level covariates. For each gene:

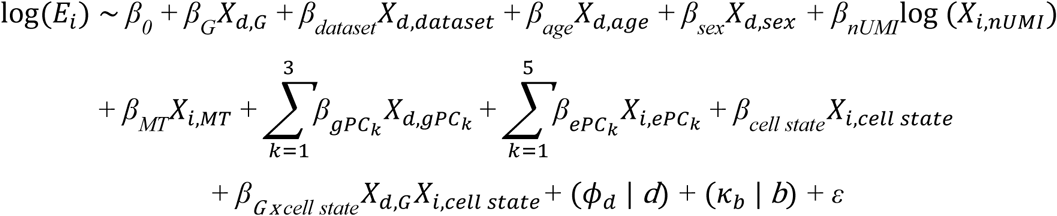

Here, *E* is the expression of the gene in cell *i*, *β_0_* is the intercept, and *ε* represents the residuals. All other *β*s represent fixed effects for various covariates in cell *i*, donor *d*, or experimental batch *b* as follows: *G* = genotype at the eQTL variant, *dataset* = dataset from which sample originates*, age* = scaled age of donor, *sex* = sex of donor, *nUMI* = number of UMI per cell (accounts for sequencing depth), *MT* = percent of mitochondrial UMIs per cell, *gPC* = genotype PCs, *ePC* = single-cell expression PCs prior to batch correction, and *cell state* = functional cell state score per cell (described above). Donor was modeled as a random individual effect (𝜙*_d_* | *d*) to account for the fact that multiple cells were sampled per individual, and experimental batch was also modeled as a random effect (𝜅*_d_* | *b*). Finally, *β_G x cell state_ X_d,G_X_i, cell state_* represents the cell state x genotype interaction term of interest.

Single-cell poisson mixed interaction models were fit using the glmer function in the lme4 R package (v 1.1-29) with the following parameters: family = “poisson”, nAGQ = 0, and control = glmerControl(optimizer = “nloptwrap”)^73^. To determine significance, we used a likelihood ratio test (LRT) comparing two models, one with and one without the cell state interaction term and calculated a p-value for the test statistic against the Chi-squared distribution with one degree of freedom. To correct for multiple hypothesis testing, we performed one permutation in which cell state scores were permuted across all cells per pathway tested, and we obtained a null LRT p-value distribution using the same framework as above with our permuted data. We then calculated q-alues for the cell state-genotype interaction term using the empirical p-value distribution across all tested eQTL using the empPvals and qvalue functions from the qvalue package (v2.16.0)^74^.

### Colocalization of GWAS and eQTL signals

Specifically for colocalization analyses, eQTL were remapped in each cell type-disease state pair with Matrix eQTL^69^ using a 1 megabase (Mb) *cis*-window, with all other modeling parameters kept constant, to broaden our search space and increase our probability of detecting colocalized variants. We assessed colocalization between our identified eQTLs in each cell type-disease state pair and the COVID-19 GWAS meta-analyses of European-ancestry subjects from the COVID-19 Host Genetics Initiative (HGI)^11^ release 7 (https://www.covid19hg.org/results/r7/). We tested two outcomes: “critical illness” and “hospitalization” (named A2 and B2, respectively by the COVID-19 HGI). A Bayesian analysis was implemented using the coloc (v5.1.0.1)^75^ R package with default settings to analyze all variants in the 1 Mb genomic locus centered on the lead eQTL in the single-cell data. We only considered GWAS loci with associations below 1 x 10^-4^. We defined colocalization as PP4 > 0.8, where PP4 corresponds to the posterior probability of colocalization between eQTL and GWAS signals. Colocalization was visualized using the R package LocusCompareR (v1.0.0)^76^ with default parameters, except for the genome parameter which was set to “hg38”. LD *r*^2^ with the lead SNP was calculated using the default “EUR” population.

---

**Fig. S1.**
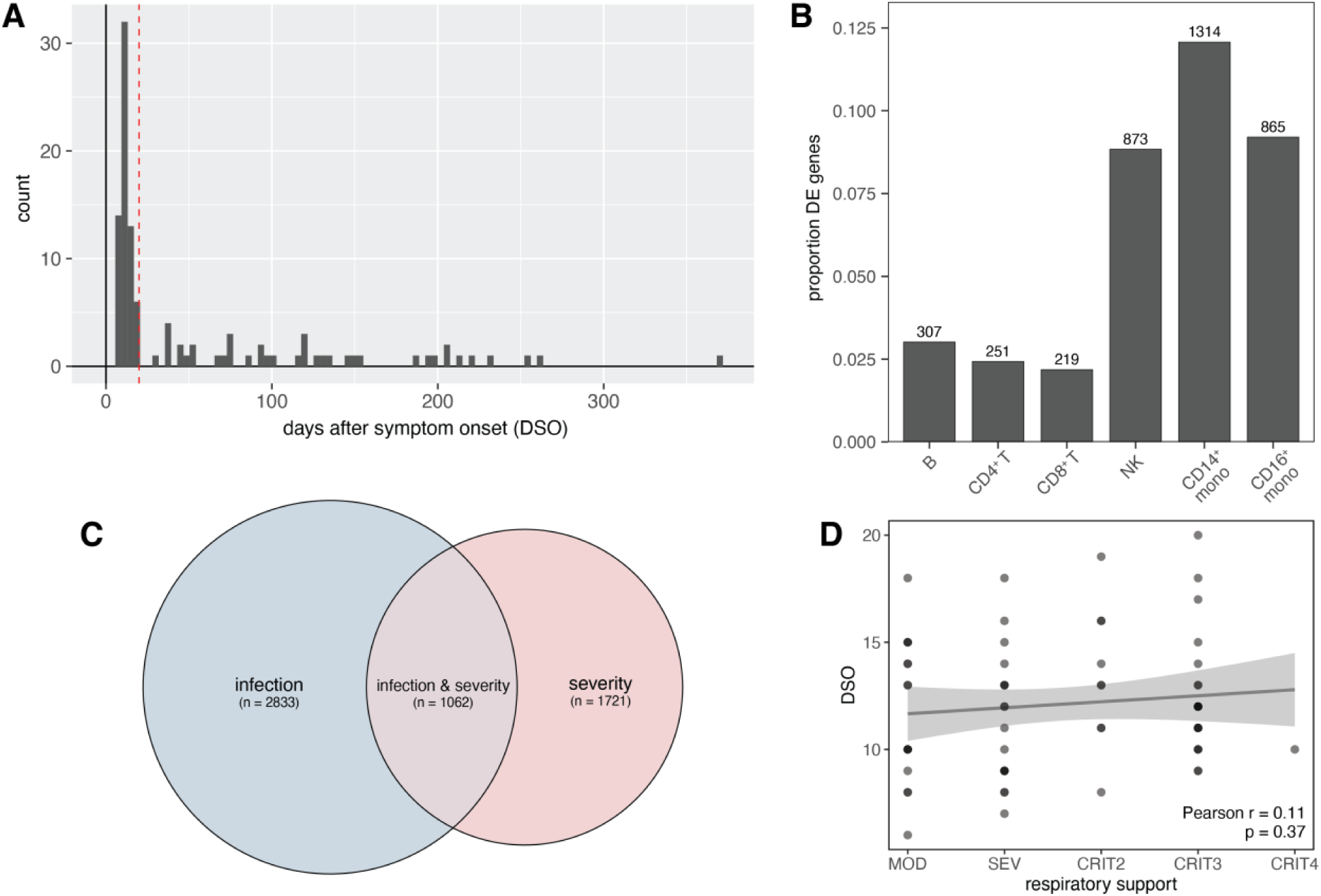
Sampling time points and global SARS-CoV-2 infection effects. **(A)** Distribution of days since symptom onset (DSO) at the time of sample collection across acute and convalescent COVID-19 patients in our cohort. Samples were considered to be in the acute phase of infection if DSO ≤ 20 (red line), and samples with DSO > 20 were considered follow-ups. **(B)** Numbers and proportions (y-axis) of genes significantly differentially expressed (|log_2_FC| > 0.5, FDR < 0.05) in COVID-19 patients compared to healthy controls. **(C)** Overlap between the set of significantly differentially expressed genes upon infection (blue circle, left) and the set of genes significantly correlated with disease severity (red circle, right). **(D)** Correlation between respiratory support score and days since symptom onset (DSO). P-value and best-fit slope were determined from a linear regression model correcting for dataset.

**Fig. S2.**
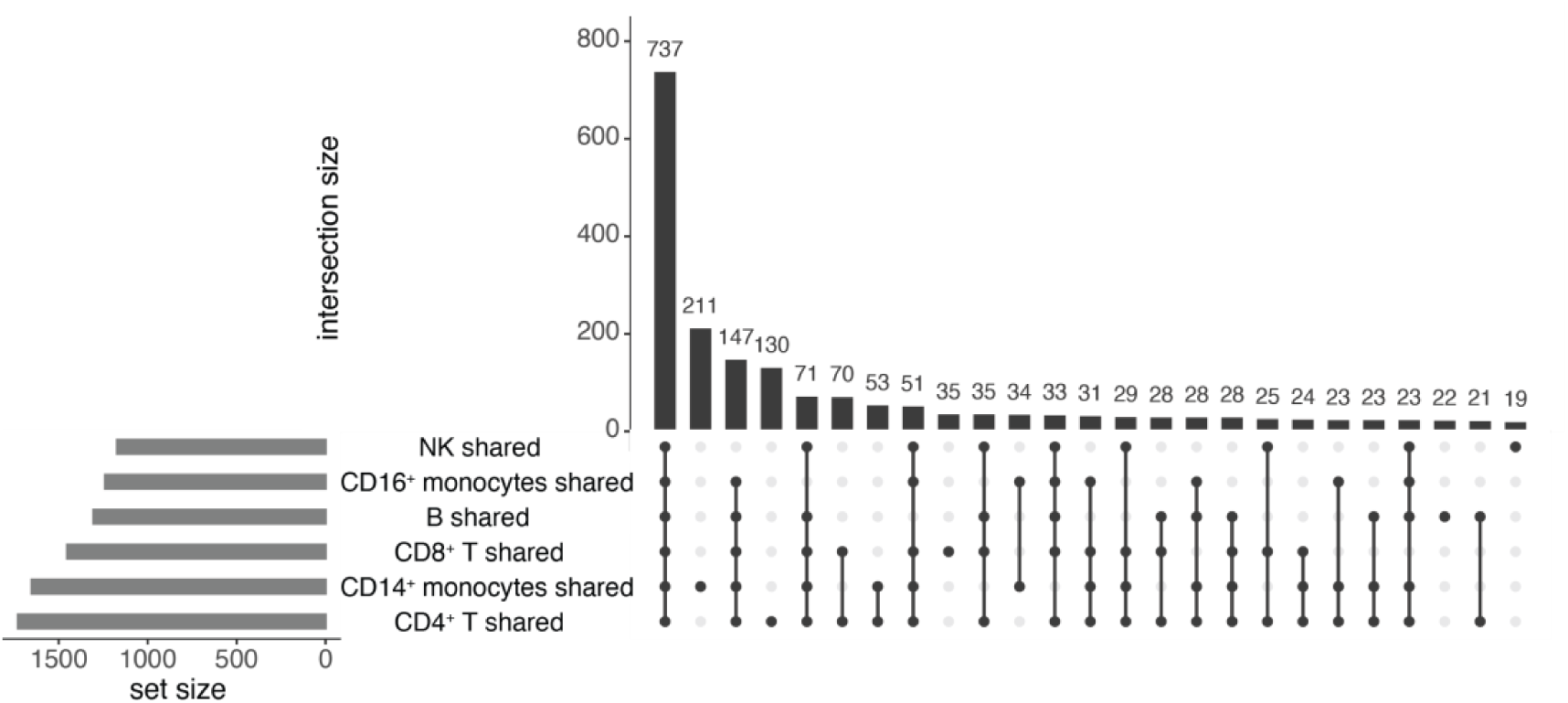
Sharing patterns among disease-state-shared eGenes. Significant eGene sharing patterns among disease-state-shared eGenes (lfsr_CTL_ < 0.1 and lfsr_COVID_ < 0.3 or lfsr_COVID_ < 0.1 and lfsr_CTL_ < 0.3) in healthy controls and COVID-19 patients across cell types.

**Fig. S3.**
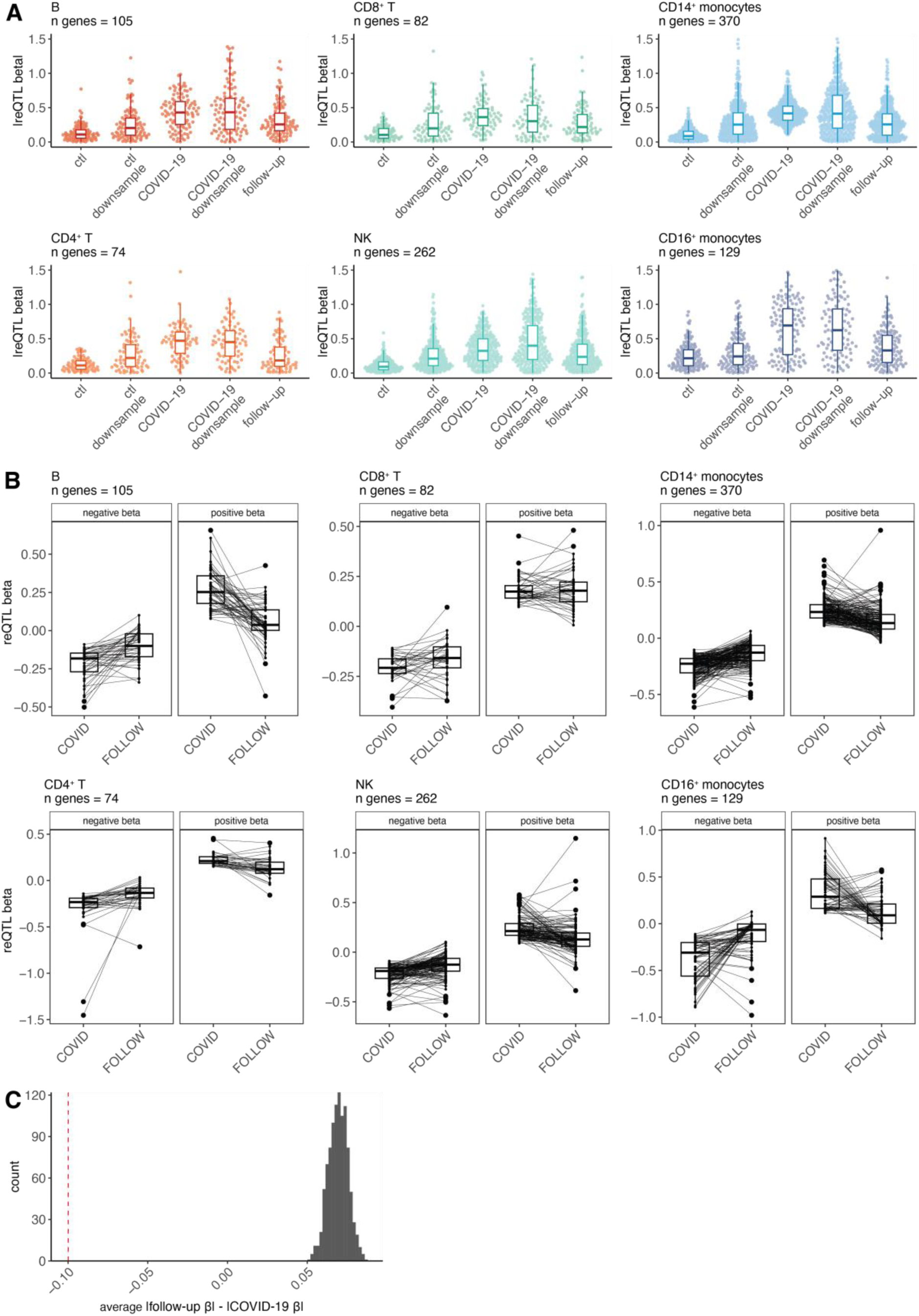
Cell type-specific response eQTL patterns. **(A)** Distribution of effect sizes for the cell type-specific reQTL sets plotted across cell types in healthy controls (“ctl”), patients (“COVID-19”), and follow-ups (“follow-up”) for the full sample set, as well as a downsampled set in the control (“ctl downsample”) and patient (“COVID-19 downsample”) groups. Downsampled sets mirrored the follow-up data structure (n = 26 samples) and were derived as follows: i) for controls, 26 individuals were randomly sampled from the control group, and ii) for patients, the 21 follow-up individuals with a corresponding acute infection time point sample were included. Here, all eQTL effect sizes are taken directly from Matrix eQTL (i.e., prior to running mash). **(B)** Paired reQTL effect sizes in COVID-19 patients (“COVID”) and follow-ups (“FOLLOW”) across cell types. The change in effect size for each gene from patient to follow-up samples is plotted as a black line. **(C)** The observed mean Δ response magnitude across the 370 CD14^+^ monocyte-specific reQTL (red dotted line) compared to the null expectation when permuting random sets of shared eGenes of the same size (n = 370) and computing their mean (n permutations = 1,000, null shown in gray). The observed mean is significantly lower (p < 0.001) than random expectation.

**Fig. S4.**
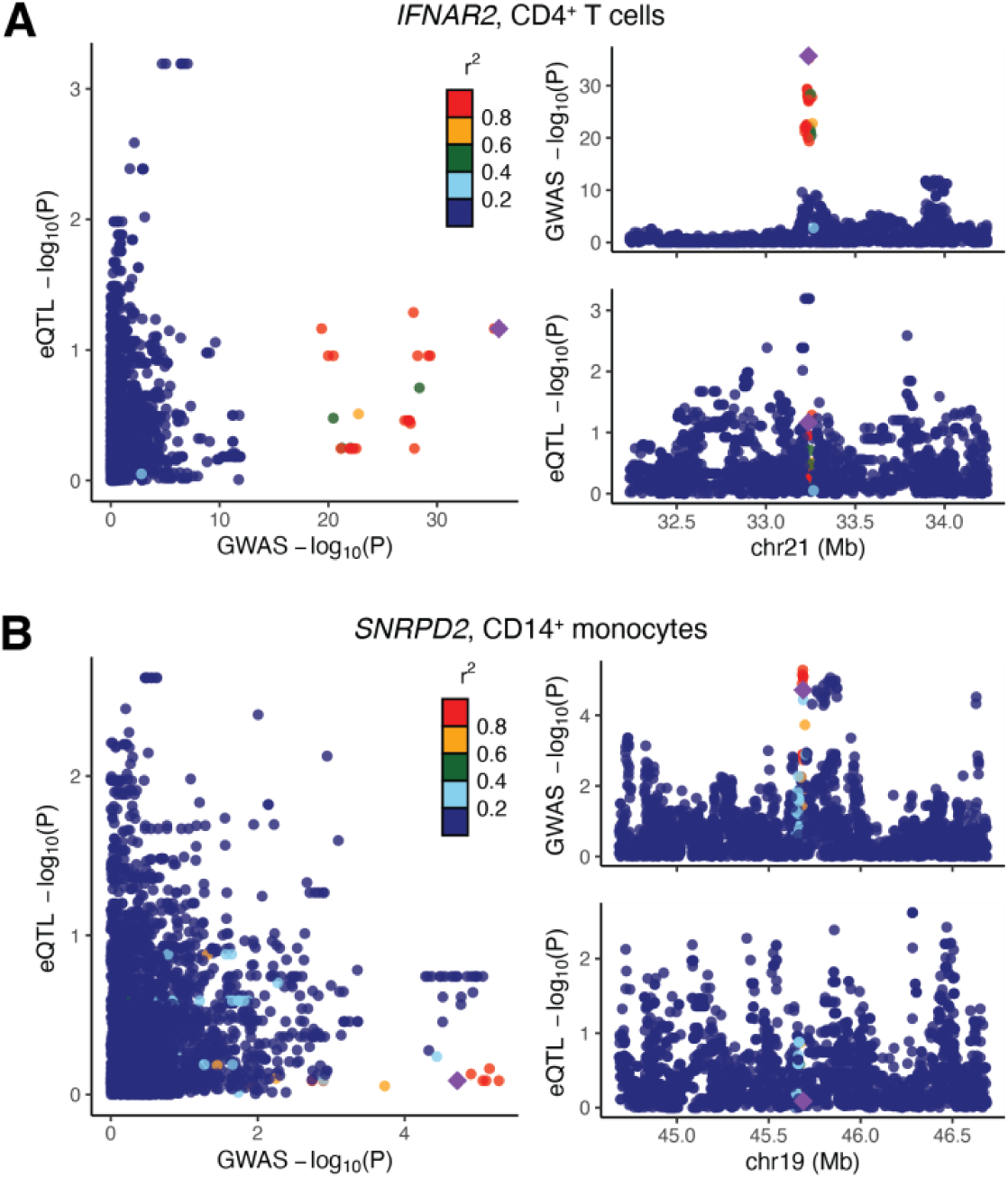
Colocalization patterns in COVID-19 follow-up samples. **(A)** The colocalization signal for the lead SNP rs9636867 (*IFNAR2*, CD4^+^ T cells, GWAS: hospitalization due to severe COVID-19) is absent in follow-ups. **(B)** The colocalization signal for the lead SNP rs7246757 (*SNRPD2*, CD14^+^ monocytes, GWAS: hospitalization due to severe COVID-19) is absent in follow-ups. For both **(A)** and **(B)**, the larger plot on the left shows the correlation between GWAS p-values (x-axis) and eQTL p-values (y-axis) in follow-ups. Smaller plots on the right show Manhattan plots for the GWAS signal (top) and the eQTL signal in follow-ups (bottom). The lead SNP is depicted as a purple diamond.

**Table S8.** Gene expression principal components (PCs) regressed in the pseudobulk eQTL analysis. PCs regressed and number of significant eQTL per cell type and disease state are reported.

| Cell type | N Regressed PCs |  |  | N genes < 0.10 FDR, Matrix eQTL |  |  |
| --- | --- | --- | --- | --- | --- | --- |
|  | *Control* | *COVID-19* | *Follow-up* | *Control* | *COVID-19* | *Follow-up* |
| *CD14<sup>+</sup>*<br>*monocytes* | 1 to 3 | 1 to 14 | 1 to 2 | 430 | 1286 | 56 |
| *CD16<sup>+</sup>*<br>*monocytes* | 1 | 1 | 1 | 10 | 49 | 6 |
| *CD4<sup>+</sup> T* | 1 to 10 | 1 to 4 | 1 to 2 | 1665 | 730 | 77 |
| *CD8<sup>+</sup> T* | 1 to 12 | 1 to 13 | 1 to 3 | 424 | 274 | 25 |
| *B* | 1 to 5 | 1 to 8 | 1 | 285 | 192 | 9 |
| *NK* | 1 to 13 | 1 to 6 | 1 to 2 | 74 | 230 | 9 |

